## Supplementary Figures and Tables for "Marker-free coselection for successive rounds of prime editing in human cells"

1. Targeting *ATP1A1* exon 17 induces cellular resistance to ouabain
2. *ATP1A1* as a selectable genomic harbor for targeted integration of transgenes
3. Targeting *ATP1A1* repeatedly for successive gene knock-in and gene tagging
4. Targeting *ATP1A1* repeatedly allows up to three sequential steps of selection
5. Coselection for adenine base editing
6. Prime editing at the *ATP1A1* locus confers resistance to ouabain
7. Robust coselection for prime editing in HeLa S3 cells
8. Coselection for HDR or PE using an EBFP to EGFP reporter system
9. *ATP1A1* as a selectable genomic harbor for targeted integration of a mTORC1 signaling reporter
10. mSc-TOSI reporter intensity in bulk coselected populations of K562 cells
11. PE3-induced indels at *MTOR* and *ATP1A1* in single cell-derived clones
12. Large PE3-mediated insertions in single cell-derived clones
13. mSc-TOSI reporter intensity in single cell-derived clones harboring *MTOR* mutations
14. Successive rounds of coselection for the isolation of double *MTOR* mutants

15. Sequential mSc-TOSI knock-in and coselection for prime editing at *MTOR* in HeLa S3 cells
16. Engineered pegRNAs (epegRNAs) with MMR-evading mutations enhance prime editing at *ATP1A1*
17. Robust coselection for prime editing at *MTOR* in U2OS cells with engineered pegRNAs
18. Characterization of large PE3-mediated insertions in bulk populations of HeLa S3 cells
19. Assessing the impact of 53BP1 inhibition on large PE-mediated insertions
20. Assessing the impact of complementary-strand nick locations on prime editing outcomes
21. Representative FACS plots for EBFP to EGFP conversion
22. Assessing the impact of complementary-strand nick locations on prime editing byproducts

**Supplementary tables**

1. Distribution of alleles in single cell-derived clones edited at *ATP1A1* via prime editing
2. Distribution of alleles in single cell-derived clones edited at *ATP1A1* and *MTOR* by coselection using engineered pegRNAs (epegRNAs) in PE2 mode.

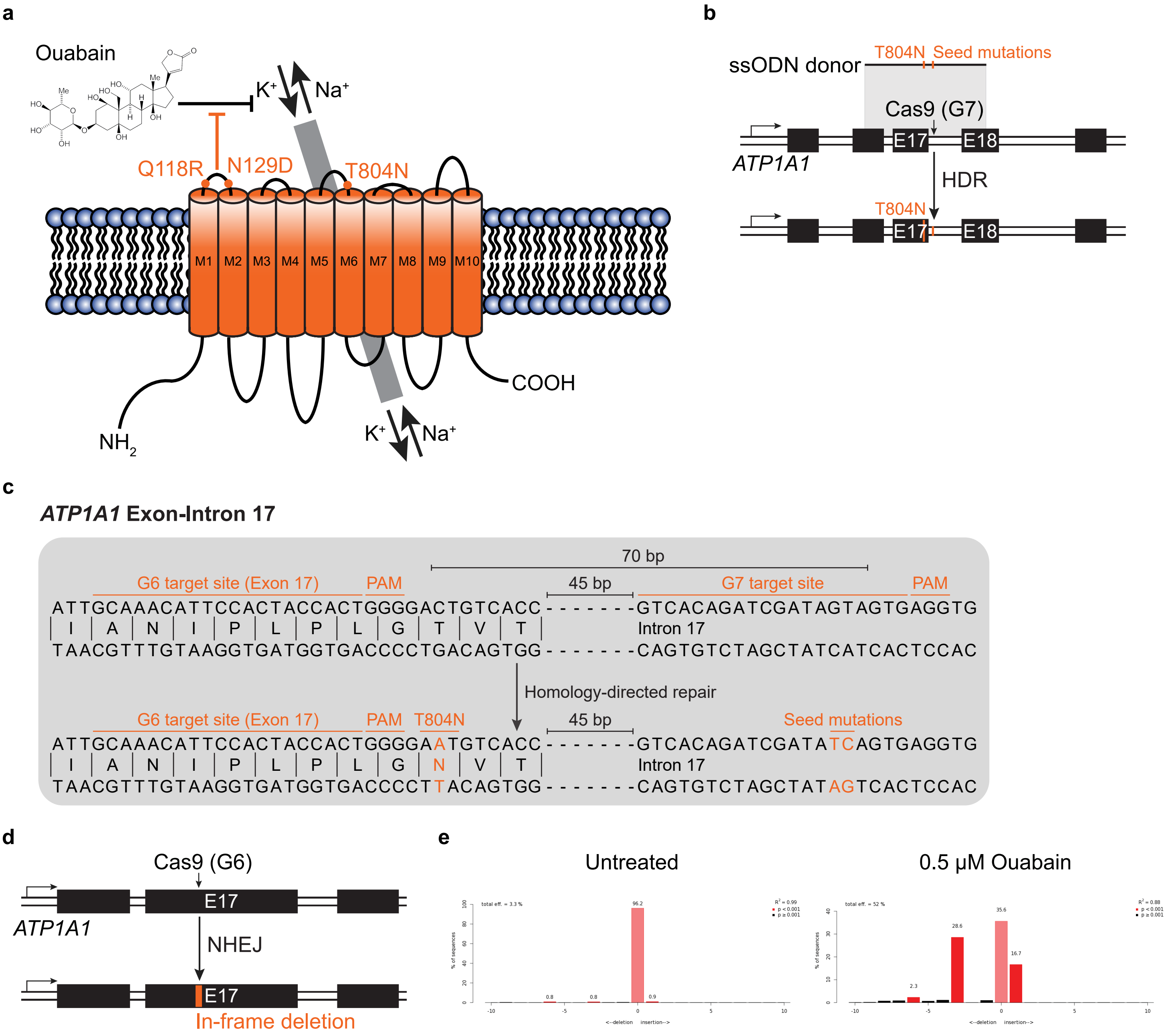

a
Ouabain
K+
Na+
Q118R
N129D
T804N
M1
M2
M3
M4
M5
M6
M7
M8
M9
M10
COOH
NH2
b
ssODN donor
T804N
Seed mutations
Cas9 (G7)
E17
E18
ATP1A1
HDR
c
ATP1A1 Exon-Intron 17
70 bp
G6 target site (Exon 17)
PAM
45 bp
G7 target site
ATTGCAAACATTCCACTACCACTGGGGACTGTCACC-------GTCACAGATCGATAGTAGTGAGGTG
I A N I P L P L G T V T
Intron 17
TAACGTTTGTAAGGTGATGGTGACCCCTGACAGTGG-------CAGTGTCTAGCTATCATCACTCCAC
Homology-directed repair
Seed mutations
ATTGCAAACATTCCACTACCACTGGGGAATGTCACC-------GTCACAGATCGATATCAGTGAGGTG
I A N I P L P L G N V T
TAACGTTTGTAAGGTGATGGTGACCCCTTACAGTGG-------CAGTGTCTAGCTATAGTCACTCCAC
d
Cas9 (G6)
NHEJ
In-frame deletion
e
Untreated
0.5 μM Ouabain

**Supplementary figure 1 (Related to Fig. 1). Targeting *ATP1A1* exon 17 induces cellular resistance to ouabain.** (**a**) Schematic representation of the $Na^+/K^+$ ATPase. The first and third extracellular loops are encoded by *ATP1A1* exon 4 and 17, respectively. (**b**) Schematic of HDR-driven editing at *ATP1A1* exon 17 using a ssODN donor. A DNA double-stranded break is introduced via SpCas9 at *ATP1A1* intron 17 and the T804N mutation is introduced via HDR at exon 17 using a ssODN donor. (**c**) Schematic representation of SpCas9 target sites used for NHEJ-driven (G6) or HDR-driven (G7) editing at *ATP1A1* exon 17 to induce cellular resistance to ouabain. The exon-intron boundary, and the distance between G7 and the T804 codon are shown. (**d**) Schematic representation of NHEJ-driven editing *ATP1A1* exon 17. (**e**) Small indels profile at *ATP1A1* exon 17 following G6-mediated cleavage and ouabain (0.5 µM) selection as determined by TIDE from Sanger sequencing. K562 cells were transfected with an eSpCas9(1.1) vector and treated with or without 0.5 µM ouabain starting 3 days post-transfection until all non-resistant cells were eliminated. $n$ = 1 experiment.

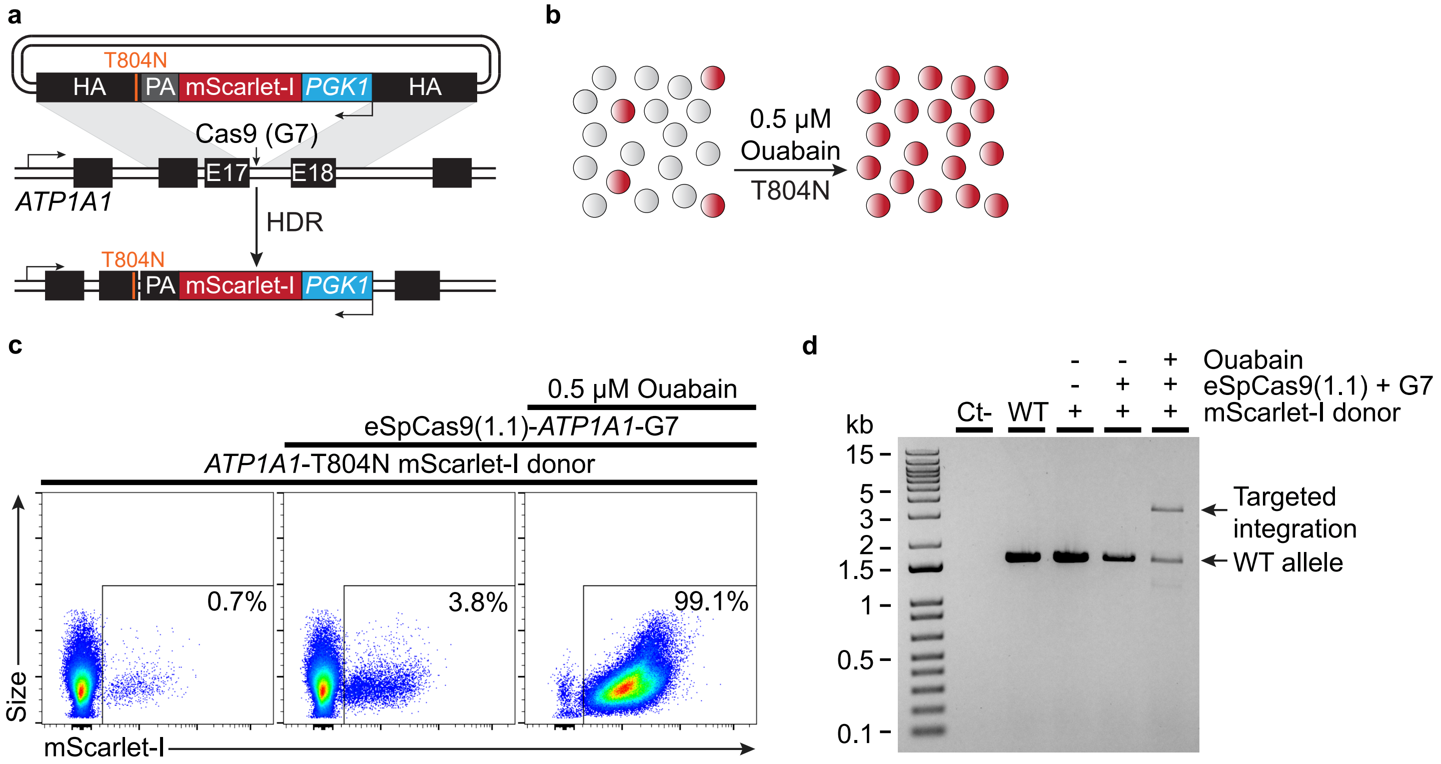

a
T804N
HA
PA
mScarlet-I
PGK1
HA
Cas9 (G7)
E17
E18
ATP1A1
HDR
T804N
PA
mScarlet-I
PGK1
b
0.5 µM
Ouabain
T804N
c
0.5 µM Ouabain
eSpCas9(1.1)-ATP1A1-G7
ATP1A1-T804N mScarlet-I donor
0.7%
3.8%
99.1%
Size
mScarlet-I
d
Ouabain
eSpCas9(1.1) + G7
mScarlet-I donor
kb
Ct-
WT
15
5
3
2
1.5
1
0.5
0.1
Targeted integration
WT allele

**Supplementary figure 2. *ATP1A1* as a selectable genomic harbor for targeted integration of transgenes.** (**a**) Schematic representation of mScarlet-I targeting to the reverse DNA strand of the *ATP1A1* locus. The T804N mutation is introduced via the left homology arm (HA) and transgene expression is driven by a human *PGK1* promoter. (**b**) Schematic of direct selection for targeted integration at *ATP1A1*. (**c**) FACS-based quantification of mScarlet-I knock-in at *ATP1A1* intron 17. K562 cells were transfected with eSpCas9(1.1) and a donor targeting *ATP1A1* and treated with or without 0.5 µM ouabain 3 days post-transfection until all non-resistant cells were eliminated. (**d**) Out-Out PCR for mScarlet-I knock-in detection at *ATP1A1* intron 17. *n* = 1 experiment.

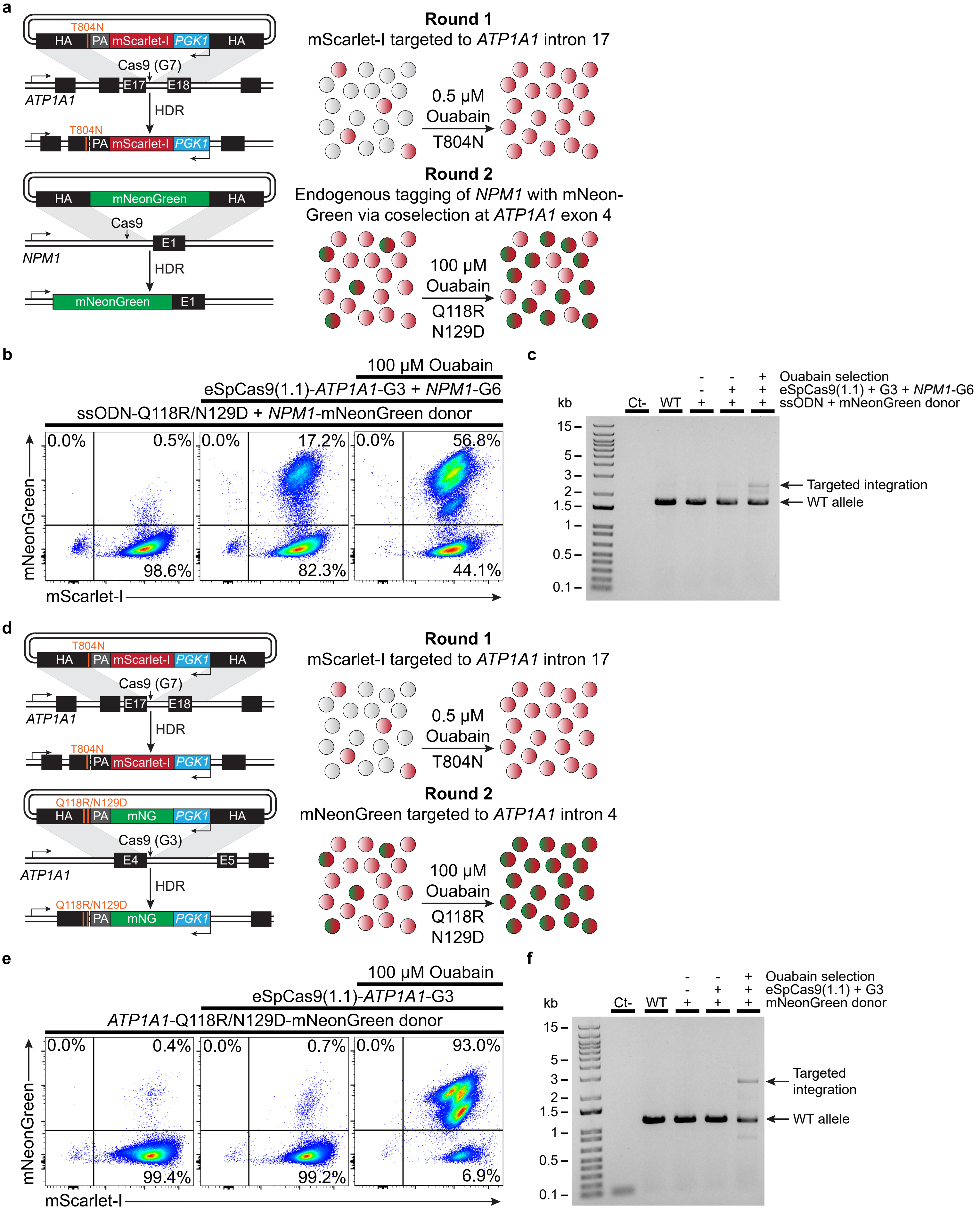
a
Round 1
mScarlet-I targeted to ATP1A1 intron 17
T804N
HA
PA
mScarlet-I
PGK1
HA
Cas9 (G7)
E17
E18
ATP1A1
HDR
0.5 µM
Ouabain
T804N
Round 2
Endogenous tagging of NPM1 with mNeon-Green via coselection at ATP1A1 exon 4
mNeonGreen
Cas9
E1
NPM1
100 µM
Ouabain
Q118R
N129D
b
100 µM Ouabain
eSpCas9(1.1)-ATP1A1-G3 + NPM1-G6
ssODN-Q118R/N129D + NPM1-mNeonGreen donor
0.0%
0.5%
98.6%
17.2%
82.3%
56.8%
44.1%
mNeonGreen
mScarlet-I
c
Ouabain selection
eSpCas9(1.1) + G3 + NPM1-G6
ssODN + mNeonGreen donor
kb
Ct-
WT
15
5
3
2
1.5
1
0.5
0.1
Targeted integration
WT allele
d
Round 1
mScarlet-I targeted to ATP1A1 intron 17
T804N
Cas9 (G7)
0.5 µM
Ouabain
Round 2
mNeonGreen targeted to ATP1A1 intron 4
Q118R/N129D
mNG
Cas9 (G3)
E4
E5
100 µM
Ouabain
Q118R
N129D
e
100 µM Ouabain
eSpCas9(1.1)-ATP1A1-G3
ATP1A1-Q118R/N129D-mNeonGreen donor
0.0%
0.4%
99.4%
0.7%
99.2%
93.0%
6.9%
mNeonGreen
mScarlet-I
f
Ouabain selection
eSpCas9(1.1) + G3
mNeonGreen donor
kb
Ct-
WT
Targeted integration
WT allele

**Supplementary figure 3. Targeting *ATP1A1* repeatedly for successive gene knock-in and gene tagging.** (**a**) Schematic representation of mScarlet-I targeting to the reverse DNA strand of the *ATP1A1* locus. The T804N mutation is introduced via the left homology arm (HA) and transgene expression is driven by a human *PGK1* promoter. Following the first selection step with a mutation that confers a medium level of resistance to ouabain (T804N), a sequential step of coselection for *NPM1* tagging with mNeonGreen is performed with a higher concentration of ouabain. (**b**) FACS-based quantification of targeted mScarlet-I integration at *ATP1A1* intron 17 and *NPM1* tagging with mNeonGreen. K562 cells stably expressing mScarlet-I (see **Supplementary Fig. 2**) were transfected with nuclease vectors and donors targeting *ATP1A1* and *NPM1* and treated with or without 100 µM ouabain 3 days post-transfection until all non-resistant cells were eliminated. (**c**) Out-Out PCR for mNeonGreen knock-in detection at *NPM1*. $n = 1$ experiment. (**d**) Same as in (**a**), but sequential gene knock-in at the *ATP1A1* locus. Following mScarlet-I knock-in at *ATP1A1* intron 17, a sequential mNeonGreen knock-in to *ATP1A1* exon 4 is performed with 100 µM ouabain. (**e**) FACS-based quantification of successive targeted integrations of mScarlet-I and mNeonGreen at *ATP1A1* intron 17 and 4, respectively. K562 cells stably expressing mScarlet-I (see **Supplementary Fig. 2**) were transfected with eSpCas9(1.1) and a donor targeting *ATP1A1* and treated with 100 µM ouabain 3 days post-transfection until all non-resistant cells were eliminated. (**f**) Out-Out PCR for mNeonGreen knock-in detection at *ATP1A1* intron 4. $n = 1$ experiment. *PGK1*, human phosphoglycerate kinase 1 promoter. PA, polyadenylation signal. HA, homology arm. E4 and E17, exon 4 and exon 17. mNG, mNeonGreen.

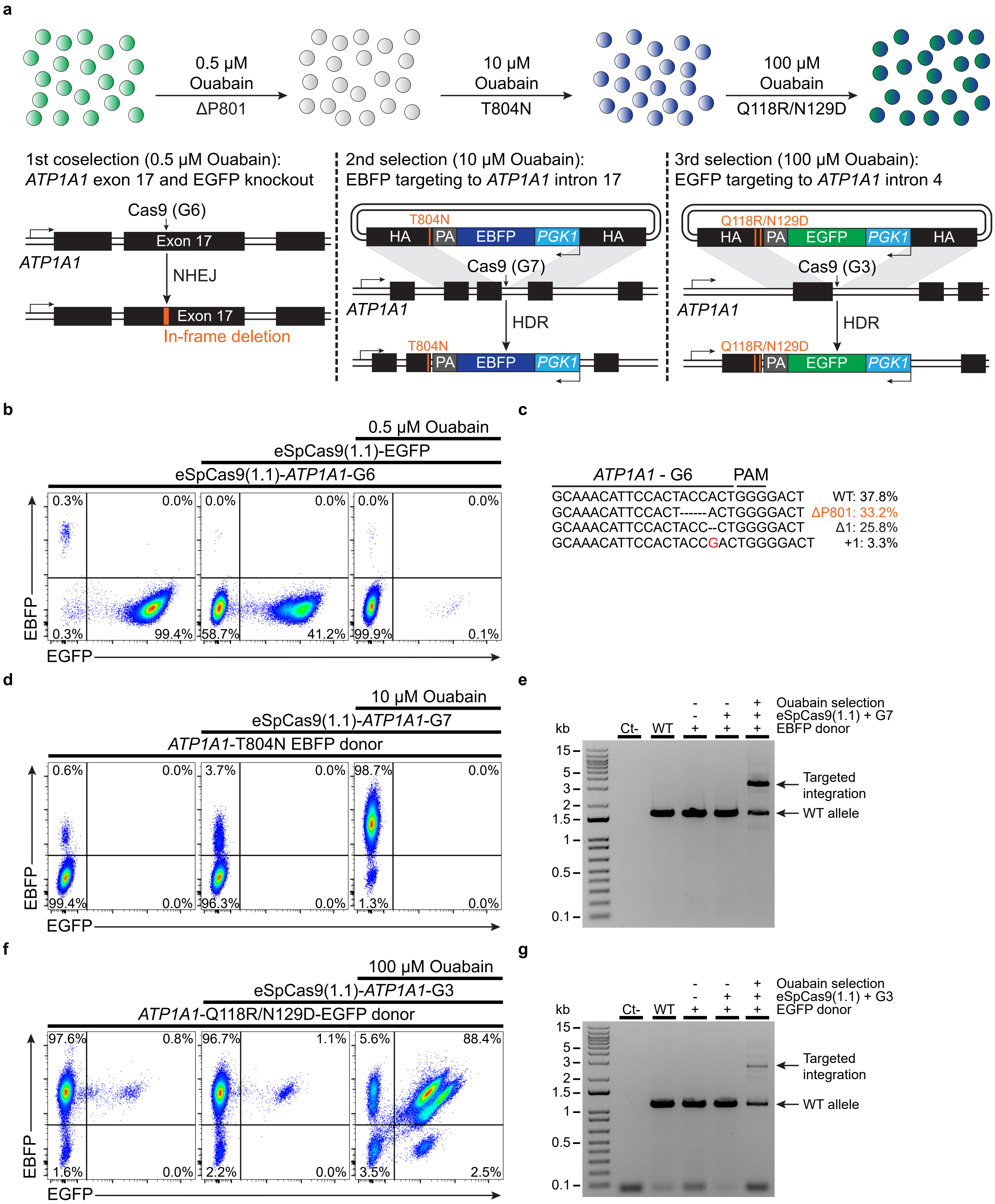

a
0.5 μM Ouabain
ΔP801
10 μM Ouabain
T804N
100 μM Ouabain
Q118R/N129D
1st coselection (0.5 μM Ouabain): ATP1A1 exon 17 and EGFP knockout
2nd selection (10 μM Ouabain): EBFP targeting to ATP1A1 intron 17
3rd selection (100 μM Ouabain): EGFP targeting to ATP1A1 intron 4
Cas9 (G6)
Exon 17
ATP1A1
NHEJ
In-frame deletion
T804N
HA
PA
EBFP
PGK1
Cas9 (G7)
HDR
Q118R/N129D
EGFP
Cas9 (G3)
b
0.5 μM Ouabain
eSpCas9(1.1)-EGFP
eSpCas9(1.1)-ATP1A1-G6
0.3% 0.0% 0.0% 0.0% 0.0% 0.0%
0.3% 99.4% 58.7% 41.2% 99.9% 0.1%
EBFP
EGFP
c
ATP1A1 - G6
PAM
GCAAACATTCCACTACCACTGGGGACT WT: 37.8%
GCAAACATTCCACT------ACTGGGGACT ΔP801: 33.2%
GCAAACATTCCACTACC--CTGGGGACT Δ1: 25.8%
GCAAACATTCCACTACCGACTGGGGACT +1: 3.3%
d
10 μM Ouabain
eSpCas9(1.1)-ATP1A1-G7
ATP1A1-T804N EBFP donor
0.6% 0.0% 3.7% 0.0% 98.7% 0.0%
99.4% 0.0% 96.3% 0.0% 1.3% 0.0%
e
kb
Ct- WT
Ouabain selection
eSpCas9(1.1) + G7
EBFP donor
15 5 3 2 1.5 1 0.5 0.1
Targeted integration
WT allele
f
100 μM Ouabain
eSpCas9(1.1)-ATP1A1-G3
ATP1A1-Q118R/N129D-EGFP donor
97.6% 0.8% 96.7% 1.1% 5.6% 88.4%
1.6% 0.0% 2.2% 0.0% 3.5% 2.5%
g
Ouabain selection
eSpCas9(1.1) + G3
EGFP donor

**Supplementary figure 4. Targeting *ATP1A1* repeatedly allows up to three sequential steps of selection.** (**a**) Schematic representation of the three steps of marker-free selection. An in-frame deletion is first introduced to *ATP1A1* exon 17 via NHEJ allowing the coselection for NHEJ events at a distant locus of interest represented by EGFP. Then, sequential targeting of expression cassettes for fluorescent proteins to *ATP1A1* introns 17 and 4 is performed by increasing the dose of ouabain at each step. (**b**) FACS-based quantification for EGFP knockout. K562 cells stably expressing EGFP from the *AAVS1* locus were transfected with nuclease vectors targeting *ATP1A1* and EGFP and treated with or without 0.5 µM ouabain 3 days post-transfection until all non-resistant cells were eliminated. (**c**) DECODR Sanger sequencing trace analysis of *ATP1A1* exon 17 following coselection for EGFP knockout. *n* = 1 experiment. (**d**) FACS-based quantification for EBFP knock-in at *ATP1A1* intron 17. Following the first step of selection, coselected K562-EGFP knockout cells were transfected with eSpCas9(1.1) and a donor to target an EBFP expression cassette to *ATP1A1* intron 17 and treated with 10 µM ouabain 3 days post-transfection until all non-resistant cells were eliminated. (**e**) Out-Out PCR for EBFP knock-in detection at *ATP1A1* intron 17. *n* = 1 experiment. (**f**) FACS-based quantification for EGFP knock-in at *ATP1A1* intron 4. Following EGFP knockout and EBFP knock-in, K562 cells were transfected with eSpCas9(1.1) and a donor to target an EGFP expression cassette to *ATP1A1* intron 4 and treated with or without 100 µM ouabain 3 days post-transfection until all non-resistant cells were eliminated. (**g**) Out-Out PCR for EGFP knock-in detection at *ATP1A1* intron 4. *n* = 1 experiment. *PGK1*, human phosphoglycerate kinase 1 promoter. PA, polyadenylation signal. HA, homology arm. E4 and E17, exon 4 and exon 17.

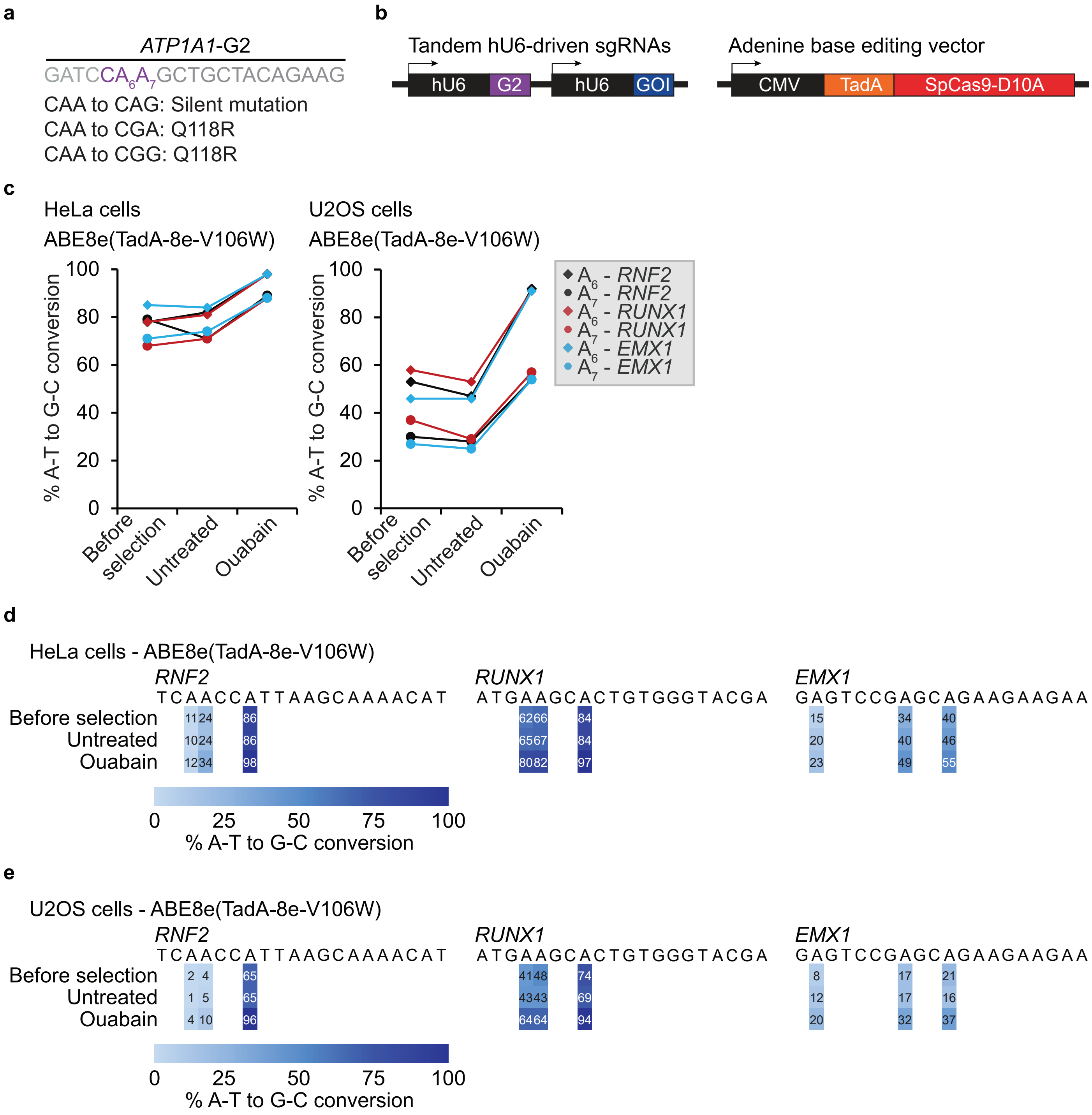
a
ATP1A1-G2
GATCCA6A7GCTGCTACAGAAG
CAA to CAG: Silent mutation
CAA to CGA: Q118R
CAA to CGG: Q118R
b
Tandem hU6-driven sgRNAs
hU6
G2
hU6
GOI
Adenine base editing vector
CMV
TadA
SpCas9-D10A
c
HeLa cells
ABE8e(TadA-8e-V106W)
U2OS cells
ABE8e(TadA-8e-V106W)
% A-T to G-C conversion
0
20
40
60
80
100
Before selection
Untreated
Ouabain
A6 - RNF2
A7 - RNF2
A6 - RUNX1
A7 - RUNX1
A6 - EMX1
A7 - EMX1
d
HeLa cells - ABE8e(TadA-8e-V106W)
RNF2
TCAACCATTAAGCAAAACAT
RUNX1
ATGAAGCACTGTGGGTACGA
EMX1
GAGTCCGAGCAGAAGAAGAA
Before selection
Untreated
Ouabain
11 24 86
10 24 86
12 34 98
62 66 84
65 67 84
80 82 97
15 34 40
20 40 46
23 49 55
0
25
50
75
100
% A-T to G-C conversion
e
U2OS cells - ABE8e(TadA-8e-V106W)
RNF2
TCAACCATTAAGCAAAACAT
RUNX1
ATGAAGCACTGTGGGTACGA
EMX1
GAGTCCGAGCAGAAGAAGAA
Before selection
Untreated
Ouabain
2 4 65
1 5 65
4 10 96
41 48 74
43 43 69
64 64 94
8 17 21
12 17 16
20 32 37
0
25
50
75
100
% A-T to G-C conversion

**Supplementary figure 5. Coselection for adenine base editing.** (**a**) Putative adenine base editing outcomes at *ATP1A1* exon 4 with SpCas9 *ATP1A1*-G2 (**b**) Schematic representation of the dual-sgRNA vector targeting *ATP1A1* and a gene of interest (GOI), and the adenine base editing vector. (**c**) Adenine base editing quantification at *ATP1A1* exon 4 (Q118R) from three experiments co-targeting *RNF2*, *RUNX1*, and *EMX1*, as determined by BEAT analysis from Sanger sequences. HeLa S3 or U2OS cells were transfected with adenine base editing and sgRNA-expressing vectors and treated with or without 0.5 µM ouabain for 10 days starting 3 days post-transfection. (**d**) Same as in (**c**) for adenine base editing at coselected GOIs in HeLa S3 cells. (**e**) Same as in (**d**) with U2OS cells. *n* = 1 experiment.

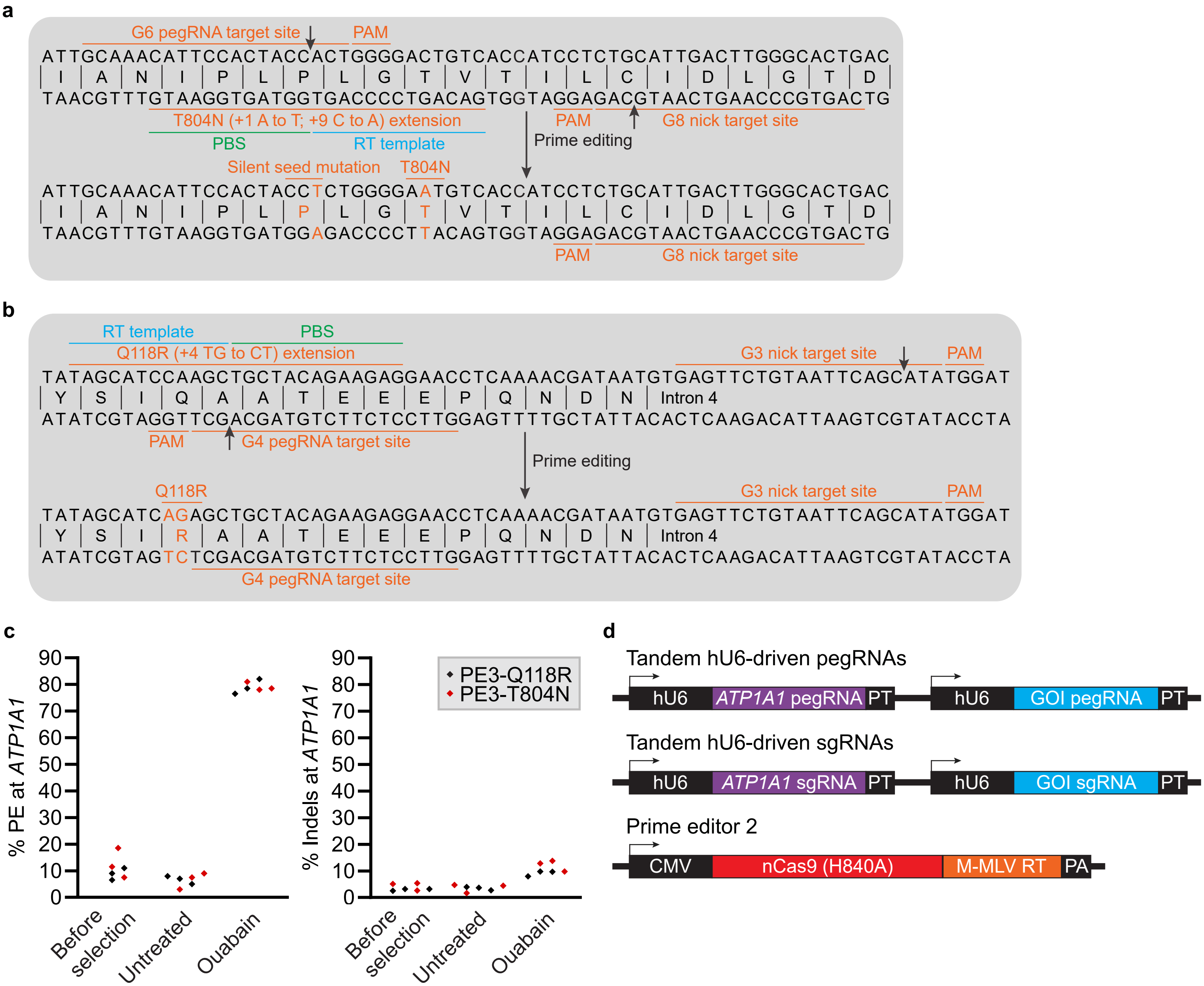
a
G6 pegRNA target site
PAM
ATTGCAAACATTCCACTACCACTGGGGACTGTCACCATCCTCTGCATTGACTTGGGCACTGAC
I A N I P L P L G T V T I L C I D L G T D
TAACGTTTGTAAGGTGATGGTGACCCCTGACAGTGGTAGGAGACGTAACTGAACCCGTGACTG
T804N (+1 A to T; +9 C to A) extension
PAM
G8 nick target site
PBS
RT template
Prime editing
Silent seed mutation
T804N
ATTGCAAACATTCCACTACCTCTGGGGAATGTCACCATCCTCTGCATTGACTTGGGCACTGAC
I A N I P L P L G T V T I L C I D L G T D
TAACGTTTGTAAGGTGATGGAGACCCCTTACAGTGGTAGGAGACGTAACTGAACCCGTGACTG
PAM
G8 nick target site
b
RT template
PBS
Q118R (+4 TG to CT) extension
G3 nick target site
PAM
TATAGCATCCAAGCTGCTACAGAAGAGGAACCTCAAAACGATAATGTGAGTTCTGTAATTCAGCATATGGAT
Y S I Q A A T E E E P Q N D N Intron 4
ATATCGTAGGTTCGACGATGTCTTCTCCTTGGAGTTTTGCTATTACACTCAAGACATTAAGTCGTATACCTA
PAM
G4 pegRNA target site
Prime editing
Q118R
G3 nick target site
PAM
TATAGCATCAGAGCTGCTACAGAAGAGGAACCTCAAAACGATAATGTGAGTTCTGTAATTCAGCATATGGAT
Y S I R A A T E E E P Q N D N Intron 4
ATATCGTAGTCTCGACGATGTCTTCTCCTTGGAGTTTTGCTATTACACTCAAGACATTAAGTCGTATACCTA
G4 pegRNA target site
c
% PE at ATP1A1
% Indels at ATP1A1
0
10
20
30
40
50
60
70
80
90
Before selection
Untreated
Ouabain
PE3-Q118R
PE3-T804N
d
Tandem hU6-driven pegRNAs
hU6
ATP1A1 pegRNA
PT
hU6
GOI pegRNA
PT
Tandem hU6-driven sgRNAs
hU6
ATP1A1 sgRNA
PT
hU6
GOI sgRNA
PT
Prime editor 2
CMV
nCas9 (H840A)
M-MLV RT
PA

**Supplementary figure 6. Prime editing at the *ATP1A1* locus confers resistance to ouabain.** (**a**) Schematic representation of the SpCas9 target sites and the reverse transcriptase (RT) template used to install the T804N mutation at *ATP1A1* exon 17 to induce cellular resistance to ouabain. (**b**) Schematic representation of the SpCas9 target sites and the reverse transcriptase (RT) template used to install the Q118R mutation at *ATP1A1* exon 4 to induce cellular resistance to ouabain. (**c**) PE and small indels quantification at *ATP1A1* as determined by BEAT and TIDE analysis from Sanger sequences. K562 cells were transfected with PE3 vectors and genomic DNA was harvested 3 days post-transfection (before selection) and cells were treated (ouabain) or not (untreated) with 0.5 µM ouabain for 10 days starting 3 days post-transfection. $n$ = 3 independent biological replicates performed at different times. (**d**) Schematic of the dual and tandem U6-driven pegRNAs and sgRNAs expression vectors along with the PE2 expression vector used to target *ATP1A1* and a gene of interest (GOI).

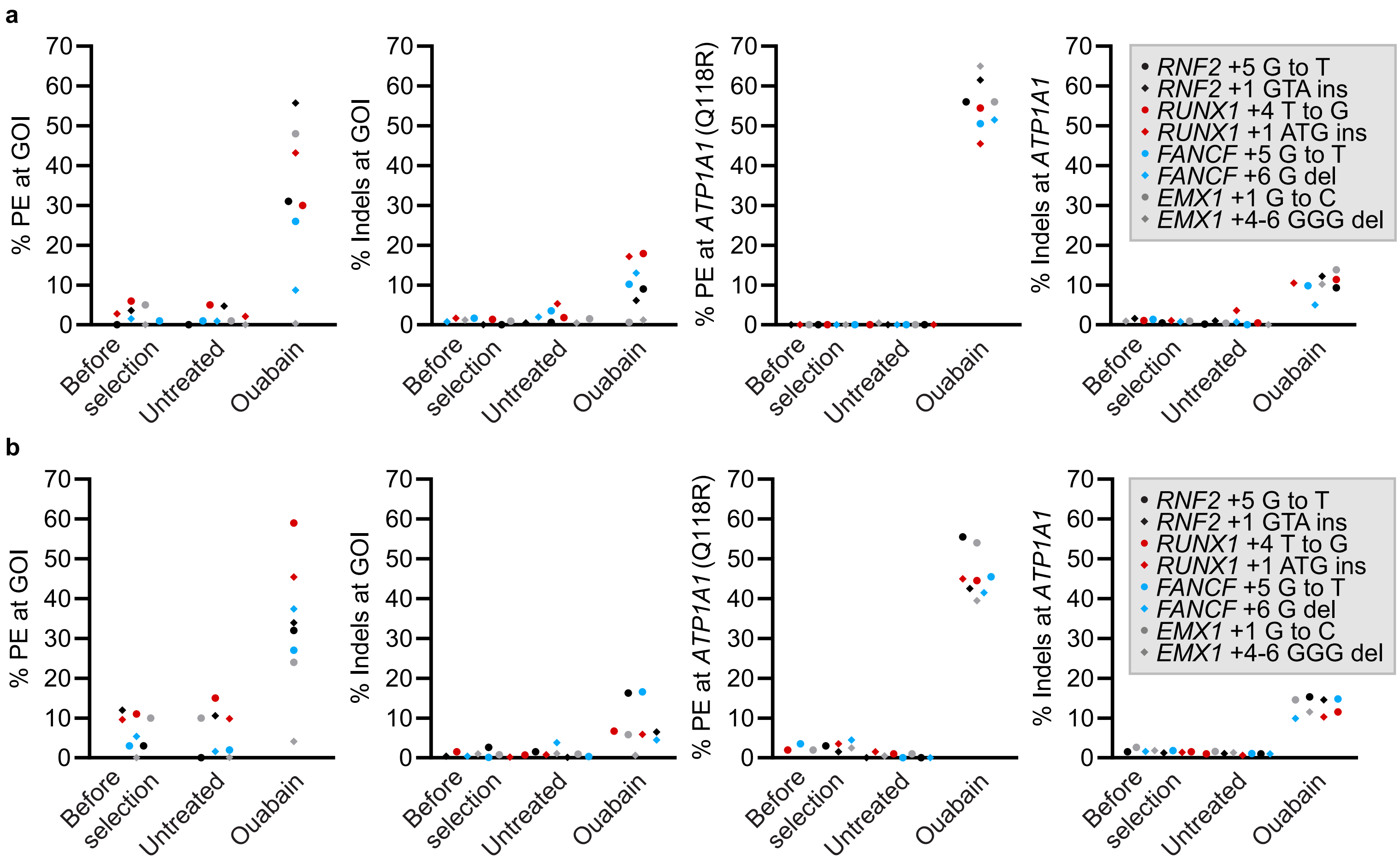

a
b
% PE at GOI
% Indels at GOI
% PE at ATP1A1 (Q118R)
% Indels at ATP1A1
0
10
20
30
40
50
60
70
Before selection
Untreated
Ouabain
RNF2 +5 G to T
RNF2 +1 GTA ins
RUNX1 +4 T to G
RUNX1 +1 ATG ins
FANCF +5 G to T
FANCF +6 G del
EMX1 +1 G to C
EMX1 +4-6 GGG del

**Supplementary figure 7 (Related to Fig. 1). Robust coselection for prime editing in HeLa S3 cells.** (**a**) PE and small indels quantification as determined by BEAT and TIDE analysis from Sanger sequences. HeLa S3 cells were transfected with PE3 vectors targeting *ATP1A1* exon 4 (Q118R) and the indicated GOI using the standard nucleofection pulse (CN-114). Genomic DNA was harvested 3 days post-transfection (before selection) and cells were treated (ouabain) or not (untreated) with 0.5 µM ouabain until all non-resistant cells were eliminated. (**b**) Same as in (**a**) using the high-efficiency nucleofection pulse (DS-150). $n$ = 2 independent biological replicates performed at different times with two different nucleofection pulses. Higher levels of editing were observed with the high-efficiency program, and we recommend using this nucleofection pulse for coselection in HeLa S3 cells.

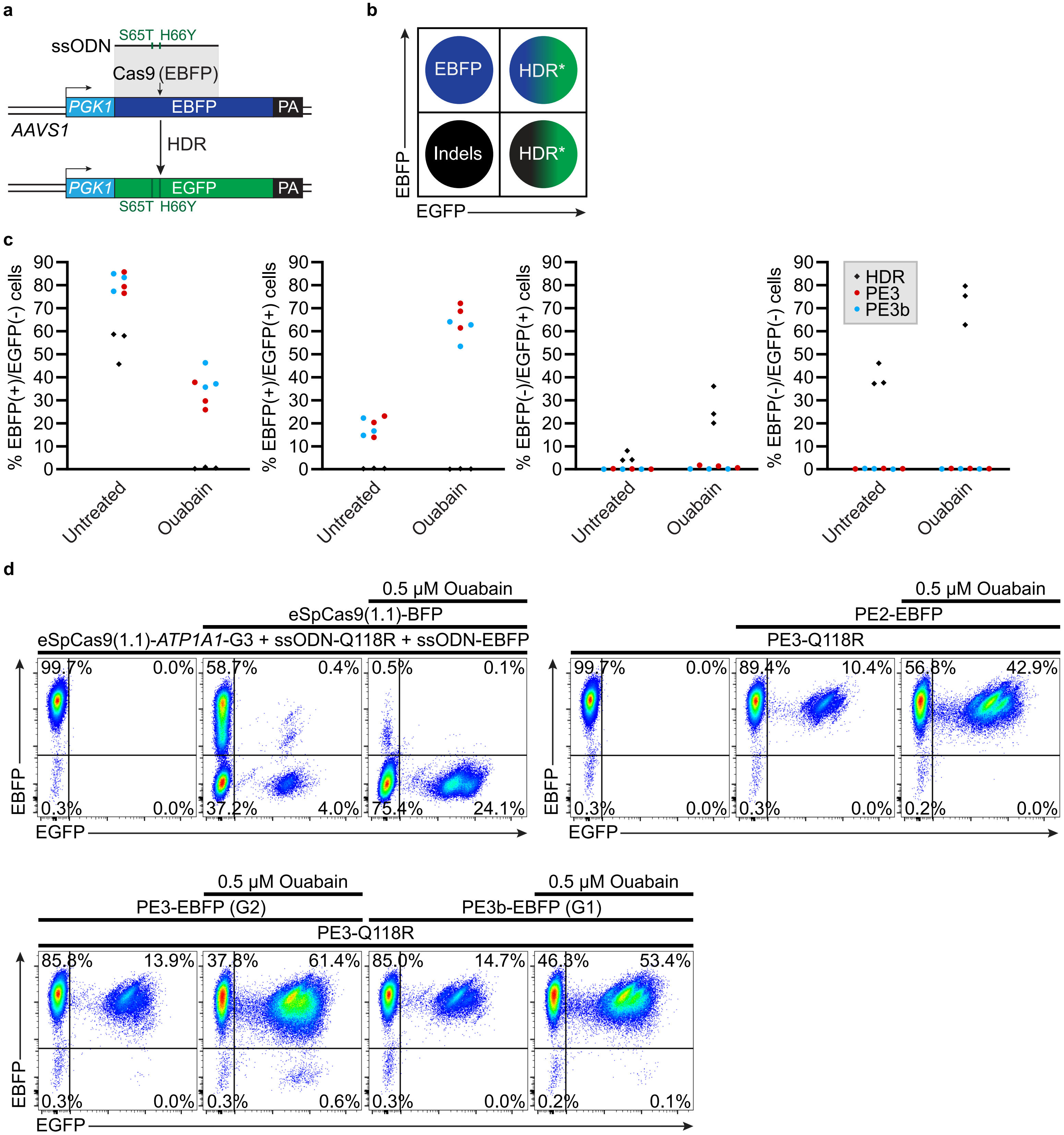

a
ssODN
S65T H66Y
Cas9 (EBFP)
PGK1
EBFP
PA
AAVS1
HDR
PGK1
EGFP
PA
S65T H66Y
b
EBFP
HDR*
Indels
HDR*
EBFP
EGFP
c
% EBFP(+)/EGFP(-) cells
% EBFP(+)/EGFP(+) cells
% EBFP(-)/EGFP(+) cells
% EBFP(-)/EGFP(-) cells
Untreated
Ouabain
HDR
PE3
PE3b
d
0.5 μM Ouabain
eSpCas9(1.1)-BFP
eSpCas9(1.1)-ATP1A1-G3 + ssODN-Q118R + ssODN-EBFP
99.7% 0.0% 0.3% 0.0%
58.7% 0.4% 37.2% 4.0%
0.5% 0.1% 75.4% 24.1%
EBFP
EGFP
0.5 μM Ouabain
PE2-EBFP
PE3-Q118R
99.7% 0.0% 0.3% 0.0%
89.4% 10.4% 0.3% 0.0%
56.8% 42.9% 0.2% 0.0%
EBFP
EGFP
0.5 μM Ouabain
PE3-EBFP (G2)
0.5 μM Ouabain
PE3b-EBFP (G1)
PE3-Q118R
85.8% 13.9% 0.3% 0.0%
37.8% 61.4% 0.3% 0.6%
85.0% 14.7% 0.3% 0.0%
46.3% 53.4% 0.2% 0.1%
EBFP
EGFP

**Supplementary figure 8. Coselection for HDR or PE using an EBFP to EGFP reporter system.** (**a**) Schematic representation of the EBFP to EGFP reporter system via HDR-based editing using a ssODN donor. (**b**) Schematic of the FACS plots following HDR-based editing of the EBFP to EGFP reporter. Precise EBFP to EGFP conversion via HDR leads to the generation of EGFP(+) cells while the fluorescence signal is lost after the introduction of indels via NHEJ-based editing. [*]Edited EGFP(+) cells can harbor small indels on remaining alleles. The EBFP transgene copy number of this cell line has not been determined. (**c**) FACS-based quantification of EBFP to EGFP conversion via HDR or PE. K562 cells stably expressing EBFP from the *AAVS1* locus were transfected with nuclease vectors and ssODN donors (HDR) or PE3/PE3b vectors targeting *ATP1A1* and EBFP. Where indicated, cells were treated with 0.5 µM ouabain starting 3 days post-transfection until all non-resistant cells were eliminated. $n = 3$ independent biological replicates performed at different times. (**d**) Representative FACS plots from the EBFP to EGFP experiments shown in (**c**). $n = 3$ independent biological replicates performed at different times with equivalent results for the HDR, PE3, and PE3b conditions, and $n = 2$ independent biological replicates for the PE2 condition.

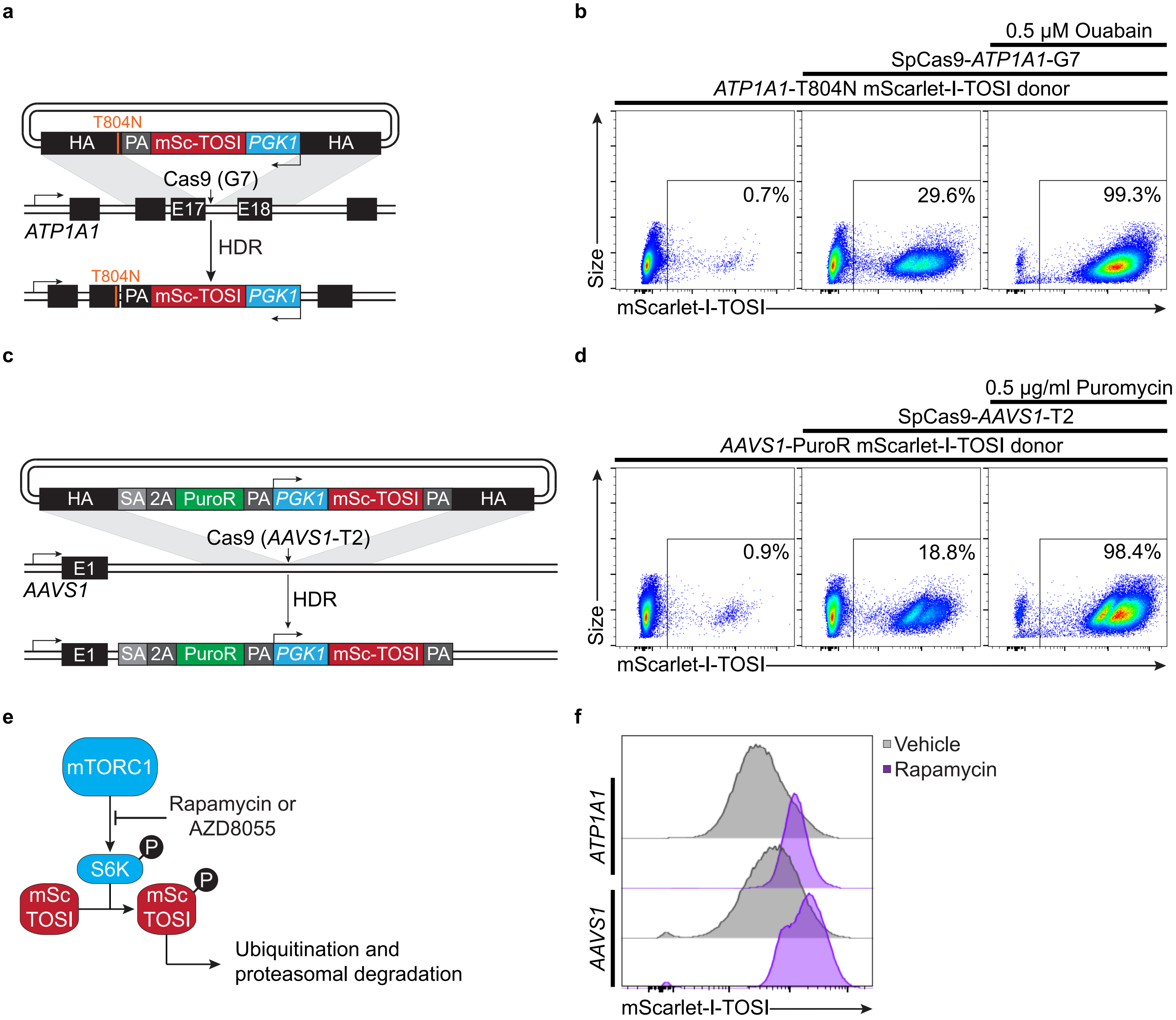
a
T804N
HA
PA
mSc-TOSI
PGK1
HA
Cas9 (G7)
E17
E18
ATP1A1
HDR
T804N
PA
mSc-TOSI
PGK1
b
0.5 µM Ouabain
SpCas9-ATP1A1-G7
ATP1A1-T804N mScarlet-I-TOSI donor
0.7%
29.6%
99.3%
Size
mScarlet-I-TOSI
c
HA
SA
2A
PuroR
PA
PGK1
mSc-TOSI
PA
HA
Cas9 (AAVS1-T2)
E1
AAVS1
HDR
E1
SA
2A
PuroR
PA
PGK1
mSc-TOSI
PA
d
0.5 µg/ml Puromycin
SpCas9-AAVS1-T2
AAVS1-PuroR mScarlet-I-TOSI donor
0.9%
18.8%
98.4%
Size
mScarlet-I-TOSI
e
mTORC1
Rapamycin or
AZD8055
S6K
P
mSc
TOSI
mSc
TOSI
P
Ubiquitination and
proteasomal degradation
f
Vehicle
Rapamycin
ATP1A1
AAVS1
mScarlet-I-TOSI

**Supplementary figure 9. *ATP1A1* as a selectable genomic harbor for targeted integration of a mTORC1 signaling reporter.** (**a**) Schematic representation of the targeted integration of mScarlet-I mTOR Signaling Indicator (mSc-TOSI) in antisense orientation within intron 17 of *ATP1A1*. The T804N mutation is introduced via the left homology arm (HA) and transgene expression is driven by the *PGK1* promoter. (**b**) FACS-based quantification of mSc-TOSI knock-in at *ATP1A1* intron 17. K562 cells were transfected with SpCas9 and the donor and treated with or without 0.5 µM ouabain starting 3 days post-transfection until all non-resistant cells were eliminated. (**c**) Schematic representation of mSc-TOSI targeting to the *AAVS1* locus. (**d**) FACS-based quantification of mSc-TOSI knock-in at *AAVS1*. K562 cells were transfected with SpCas9 and a donor targeting *AAVS1* and treated with or without 0.5 µg/ml puromycin starting 3 days post-transfection until all non-resistant cells were eliminated. (**e**) Schematic of mSc-TOSI degradation under mTORC1 signaling. (**f**) mSc-TOSI fluorescence intensity after stable integration at the *ATP1A1* and *AAVS1* loci in presence and absence of rapamycin. Where indicated, cells were treated with 500 nM rapamycin or vehicle control for 24 hours before FACS analysis. $n$ = 1 experiment. SA, splicing acceptor. *PGK1*, human phosphoglycerate kinase 1 promoter. PA, polyadenylation signal. HA, homology arm. E, exon.

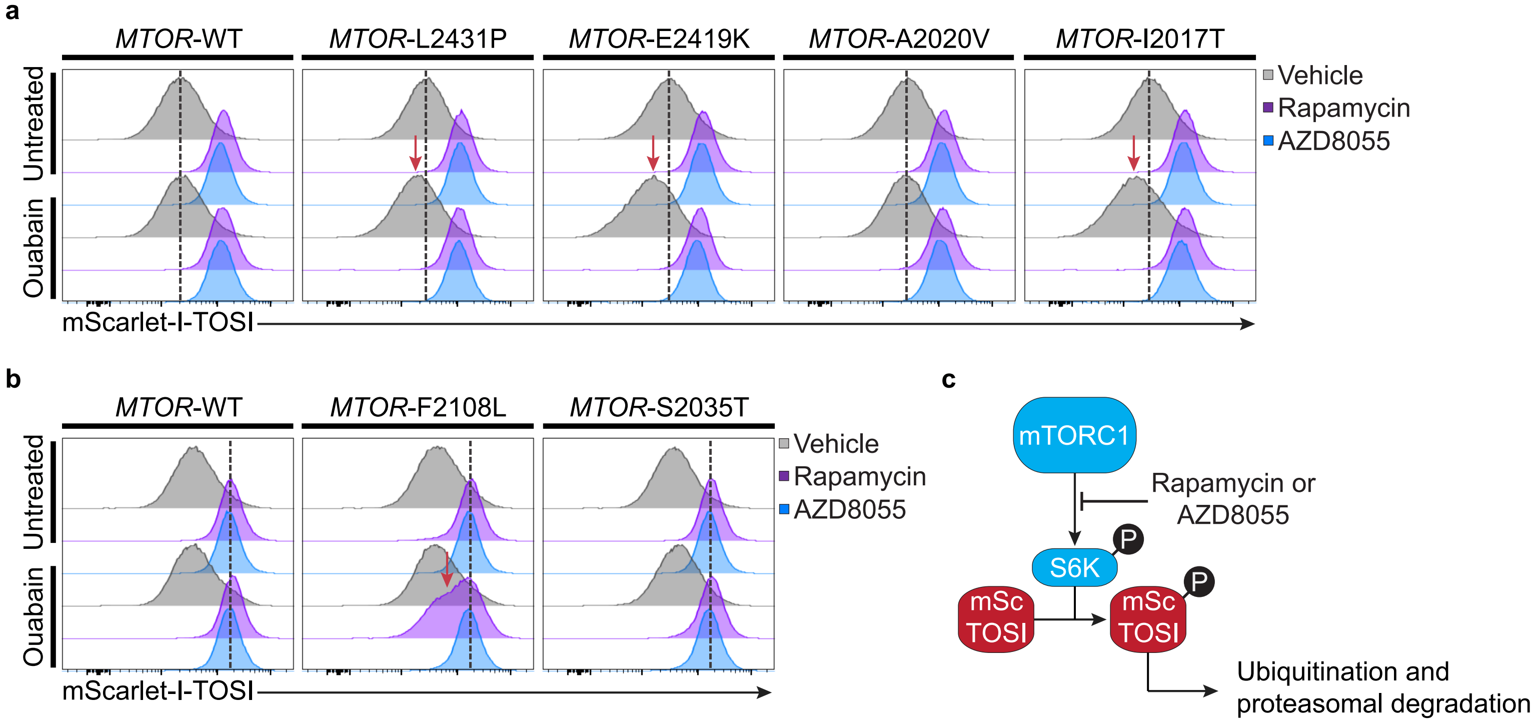

a
MTOR-WT
MTOR-L2431P
MTOR-E2419K
MTOR-A2020V
MTOR-I2017T
Untreated
Ouabain
Vehicle
Rapamycin
AZD8055
mScarlet-I-TOSI
b
MTOR-WT
MTOR-F2108L
MTOR-S2035T
Untreated
Ouabain
Vehicle
Rapamycin
AZD8055
mScarlet-I-TOSI
c
mTORC1
Rapamycin or AZD8055
S6K
P
mSc TOSI
mSc TOSI
P
Ubiquitination and proteasomal degradation

**Supplementary figure 10 (Related to Fig. 3). mSc-TOSI reporter intensity in bulk coselected populations of K562 cells.** (**a**) Histogram plot of mSc-TOSI intensity after coselection for *MTOR* hyperactivating mutations. As described in **Fig. 3**, K562 cells stably expressing the mSc-TOSI reporter were transfected with PE3 vectors targeting *ATP1A1* exon 4 (Q118R) and *MTOR*. Cells were treated with or without 100 µM ouabain 3 days post-transfection until all non-resistant cells were eliminated. After coselection, cells were treated for 24 hours with 50 nM rapamycin or 50 nM AZD8055 before FACS analysis. (**b**) Same as in (**a**) with *MTOR* rapamycin-resistant mutations. Representative images are from one of two independent biological replicates performed at different times with equivalent results. (**c**) Schematic of mSc-TOSI degradation under mTORC1 signaling.

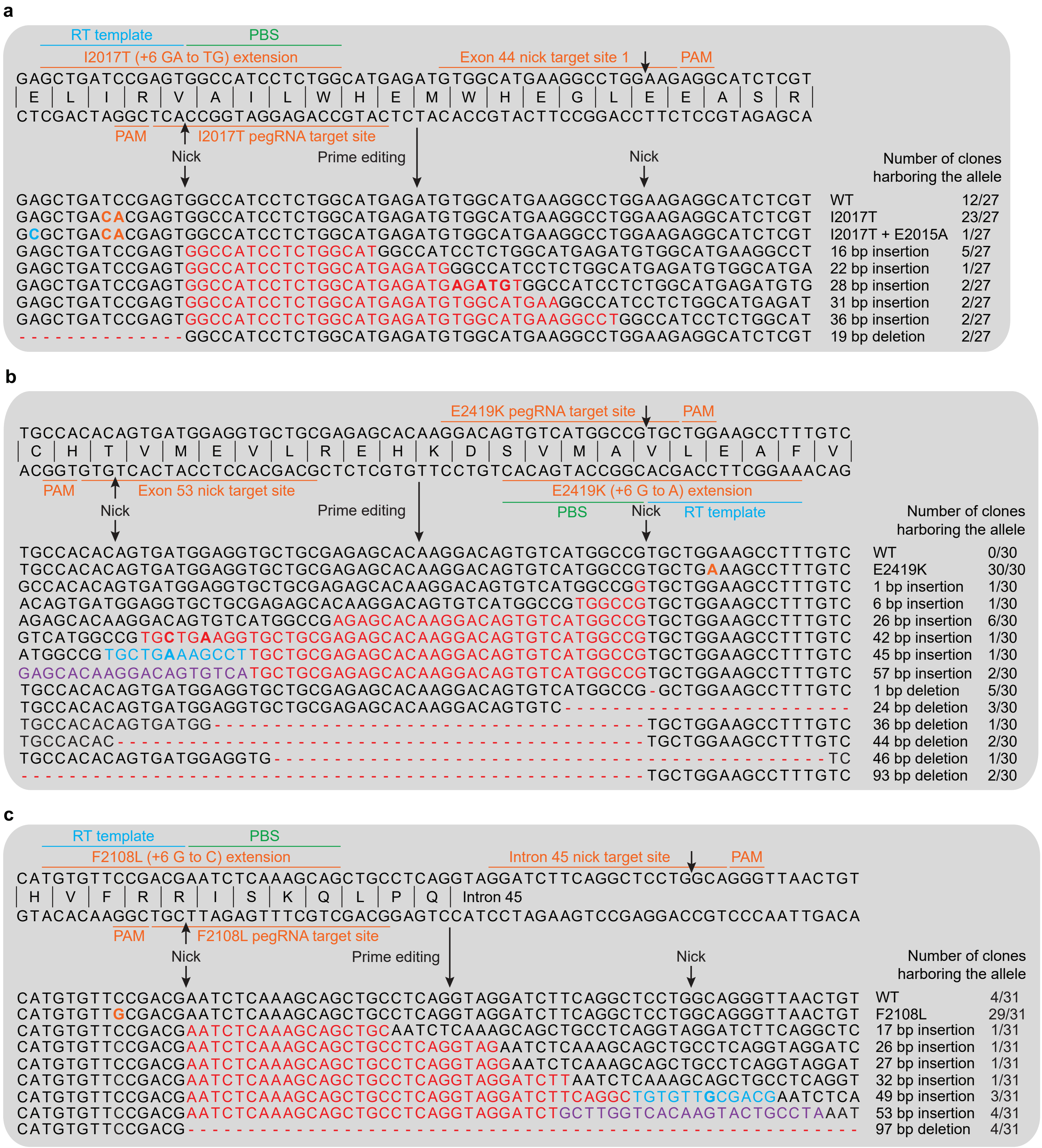
a
RT template
PBS
I2017T (+6 GA to TG) extension
Exon 44 nick target site 1
PAM
GAGCTGATCCGAGTGGCCATCCTCTGGCATGAGATGTGGCATGAAGGCCTGGAAGAGGCATCTCGT
E L I R V A I L W H E M W H E G L E E A S R
CTCGACTAGGCTCACCGGTAGGAGACCGTACTCTACACCGTACTTCCGGACCTTCTCCGTAGAGCA
PAM
I2017T pegRNA target site
Nick
Prime editing
Nick
Number of clones harboring the allele
GAGCTGATCCGAGTGGCCATCCTCTGGCATGAGATGTGGCATGAAGGCCTGGAAGAGGCATCTCGT WT 12/27
GAGCTGACACGAGTGGCCATCCTCTGGCATGAGATGTGGCATGAAGGCCTGGAAGAGGCATCTCGT I2017T 23/27
GCGCTGACACGAGTGGCCATCCTCTGGCATGAGATGTGGCATGAAGGCCTGGAAGAGGCATCTCGT I2017T + E2015A 1/27
GAGCTGATCCGAGTGGCCATCCTCTGGCATGGCCATCCTCTGGCATGAGATGTGGCATGAAGGCCT 16 bp insertion 5/27
GAGCTGATCCGAGTGGCCATCCTCTGGCATGAGATGGCCATCCTCTGGCATGAGATGTGGCATGA 22 bp insertion 1/27
GAGCTGATCCGAGTGGCCATCCTCTGGCATGAGATGAGATGTGGCCATCCTCTGGCATGAGATGTG 28 bp insertion 2/27
GAGCTGATCCGAGTGGCCATCCTCTGGCATGAGATGTGGCATGAAGGCCATCCTCTGGCATGAGAT 31 bp insertion 2/27
GAGCTGATCCGAGTGGCCATCCTCTGGCATGAGATGTGGCATGAAGGCCTGGCCATCCTCTGGCAT 36 bp insertion 2/27
---------------GGCCATCCTCTGGCATGAGATGTGGCATGAAGGCCTGGAAGAGGCATCTCGT 19 bp deletion 2/27
b
E2419K pegRNA target site
PAM
TGCCACACAGTGATGGAGGTGCTGCGAGAGCACAAGGACAGTGTCATGGCCGTGCTGGAAGCCTTTGTC
C H T V M E V L R E H K D S V M A V L E A F V
ACGGTGTGTCACTACCTCCACGACGCTCTCGTGTTCCTGTCACAGTACCGGCACGACCTTCGGAAACAG
PAM
Exon 53 nick target site
E2419K (+6 G to A) extension
PBS
RT template
Nick
Prime editing
Nick
Number of clones harboring the allele
TGCCACACAGTGATGGAGGTGCTGCGAGAGCACAAGGACAGTGTCATGGCCGTGCTGGAAGCCTTTGTC WT 0/30
TGCCACACAGTGATGGAGGTGCTGCGAGAGCACAAGGACAGTGTCATGGCCGTGCTGAAAGCCTTTGTC E2419K 30/30
GCCACACAGTGATGGAGGTGCTGCGAGAGCACAAGGACAGTGTCATGGCCGGTGCTGGAAGCCTTTGTC 1 bp insertion 1/30
ACAGTGATGGAGGTGCTGCGAGAGCACAAGGACAGTGTCATGGCCGTGGCCGTGCTGGAAGCCTTTGTC 6 bp insertion 1/30
AGAGCACAAGGACAGTGTCATGGCCGAGAGCACAAGGACAGTGTCATGGCCGTGCTGGAAGCCTTTGTC 26 bp insertion 6/30
GTCATGGCCGTGCTGAAGGTGCTGCGAGAGCACAAGGACAGTGTCATGGCCGTGCTGGAAGCCTTTGTC 42 bp insertion 1/30
ATGGCCGTGCTGAAAGCCTTGCTGCGAGAGCACAAGGACAGTGTCATGGCCGTGCTGGAAGCCTTTGTC 45 bp insertion 1/30
GAGCACAAGGACAGTGTCATGCTGCGAGAGCACAAGGACAGTGTCATGGCCGTGCTGGAAGCCTTTGTC 57 bp insertion 2/30
TGCCACACAGTGATGGAGGTGCTGCGAGAGCACAAGGACAGTGTCATGGCCG-GCTGGAAGCCTTTGTC 1 bp deletion 5/30
TGCCACACAGTGATGGAGGTGCTGCGAGAGCACAAGGACAGTGTC------------------------ 24 bp deletion 3/30
TGCCACACAGTGATGG------------------------------------TGCTGGAAGCCTTTGTC 36 bp deletion 1/30
TGCCACAC--------------------------------------------TGCTGGAAGCCTTTGTC 44 bp deletion 2/30
TGCCACACAGTGATGGAGGTG----------------------------------------------TC 46 bp deletion 1/30
----------------------------------------------------TGCTGGAAGCCTTTGTC 93 bp deletion 2/30
c
RT template
PBS
F2108L (+6 G to C) extension
Intron 45 nick target site
PAM
CATGTGTTCCGACGAATCTCAAAGCAGCTGCCTCAGGTAGGATCTTCAGGCTCCTGGCAGGGTTAACTGT
H V F R R I S K Q L P Q Intron 45
GTACACAAGGCTGCTTAGAGTTTCGTCGACGGAGTCCATCCTAGAAGTCCGAGGACCGTCCCAATTGACA
PAM
F2108L pegRNA target site
Nick
Prime editing
Nick
Number of clones harboring the allele
CATGTGTTCCGACGAATCTCAAAGCAGCTGCCTCAGGTAGGATCTTCAGGCTCCTGGCAGGGTTAACTGT WT 4/31
CATGTGTTGCGACGAATCTCAAAGCAGCTGCCTCAGGTAGGATCTTCAGGCTCCTGGCAGGGTTAACTGT F2108L 29/31
CATGTGTTCCGACGAATCTCAAAGCAGCTGCAATCTCAAAGCAGCTGCCTCAGGTAGGATCTTCAGGCTC 17 bp insertion 1/31
CATGTGTTCCGACGAATCTCAAAGCAGCTGCCTCAGGTAGAATCTCAAAGCAGCTGCCTCAGGTAGGATC 26 bp insertion 1/31
CATGTGTTCCGACGAATCTCAAAGCAGCTGCCTCAGGTAGGAATCTCAAAGCAGCTGCCTCAGGTAGGAT 27 bp insertion 1/31
CATGTGTTCCGACGAATCTCAAAGCAGCTGCCTCAGGTAGGATCTTAATCTCAAAGCAGCTGCCTCAGGT 32 bp insertion 1/31
CATGTGTTCCGACGAATCTCAAAGCAGCTGCCTCAGGTAGGATCTTCAGGCTGTGTTGCGACGAATCTCA 49 bp insertion 3/31
CATGTGTTCCGACGAATCTCAAAGCAGCTGCCTCAGGTAGGATCTGCTTGGTCACAAGTACTGCCTAAAT 53 bp insertion 4/31
CATGTGTTCCGACG-------------------------------------------------------- 97 bp deletion 4/31

**d**

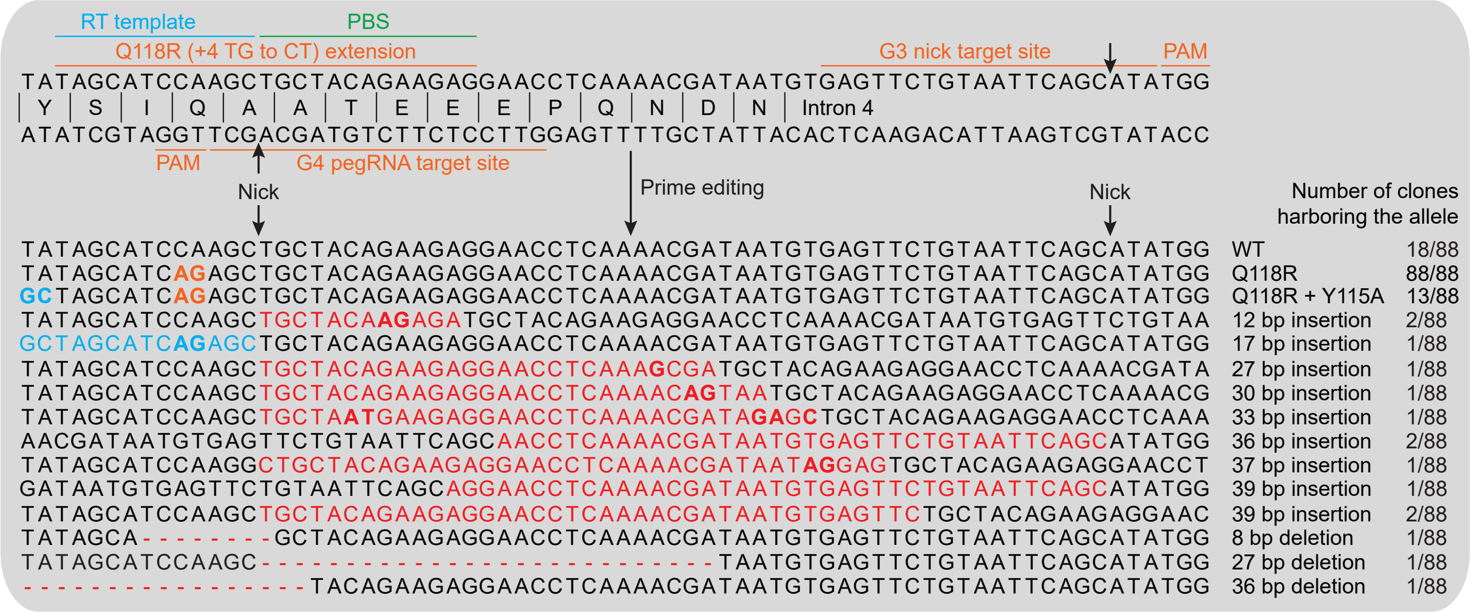

RT template
PBS
Q118R (+4 TG to CT) extension
G3 nick target site
PAM
TATAGCATCCAAGCTGCTACAGAAGAGGAACCTCAAAACGATAATGTGAGTTCTGTAATTCAGCATATGG
Y S I Q A A T E E E P Q N D N Intron 4
ATATCGTAGGTTCGACGATGTCTTCTCCTTGGAGTTTTGCTATTACACTCAAGACATTAAGTCGTATACC
PAM
G4 pegRNA target site
Nick
Prime editing
Nick
Number of clones harboring the allele
TATAGCATCCAAGCTGCTACAGAAGAGGAACCTCAAAACGATAATGTGAGTTCTGTAATTCAGCATATGG WT 18/88
TATAGCATCAGAGCTGCTACAGAAGAGGAACCTCAAAACGATAATGTGAGTTCTGTAATTCAGCATATGG Q118R 88/88
GCTAGCATCAGAGCTGCTACAGAAGAGGAACCTCAAAACGATAATGTGAGTTCTGTAATTCAGCATATGG Q118R + Y115A 13/88
TATAGCATCCAAGCTGCTACAAGAGATGCTACAGAAGAGGAACCTCAAAACGATAATGTGAGTTCTGTAA 12 bp insertion 2/88
GCTAGCATCAGAGCTGCTACAGAAGAGGAACCTCAAAACGATAATGTGAGTTCTGTAATTCAGCATATGG 17 bp insertion 1/88
TATAGCATCCAAGCTGCTACAGAAGAGGAACCTCAAAGCGATGCTACAGAAGAGGAACCTCAAAACGATA 27 bp insertion 1/88
TATAGCATCCAAGCTGCTACAGAAGAGGAACCTCAAAACAGTAATGCTACAGAAGAGGAACCTCAAAACG 30 bp insertion 1/88
TATAGCATCCAAGCTGCTAATGAAGAGGAACCTCAAAACGATAGAGCTGCTACAGAAGAGGAACCTCAAA 33 bp insertion 1/88
AACGATAATGTGAGTTCTGTAATTCAGCAACCTCAAAACGATAATGTGAGTTCTGTAATTCAGCATATGG 36 bp insertion 2/88
TATAGCATCCAAGGCTGCTACAGAAGAGGAACCTCAAAACGATAATAGGAGTGCTACAGAAGAGGAACCT 37 bp insertion 1/88
GATAATGTGAGTTCTGTAATTCAGCAGGAACCTCAAAACGATAATGTGAGTTCTGTAATTCAGCATATGG 39 bp insertion 1/88
TATAGCATCCAAGCTGCTACAGAAGAGGAACCTCAAAACGATAATGTGAGTTCTGCTACAGAAGAGGAAC 39 bp insertion 2/88
TATAGCA---------GCTACAGAAGAGGAACCTCAAAACGATAATGTGAGTTCTGTAATTCAGCATATGG 8 bp deletion 1/88
TATAGCATCCAAGC---------------------------TAATGTGAGTTCTGTAATTCAGCATATGG 27 bp deletion 1/88
-----------------TACAGAAGAGGAACCTCAAAACGATAATGTGAGTTCTGTAATTCAGCATATGG 36 bp deletion 1/88

**Supplementary figure 11 (Related to Fig. 3 and Table 1). PE3-induced indels at *MTOR* and *ATP1A1* in single cell-derived clones.** (**a**) Schematic representation of the targeting strategy and the PE-induced insertions and deletions at *MTOR* exon 44 as determined by DECODR Sanger sequencing trace analysis. Single cell-derived K562 clones were isolated in methylcellulose-based semi-solid RPMI media supplemented with 100 µM ouabain and genomic DNA was harvested after coselection. (**b**) Same as in (**a**) for *MTOR* exon 53. (**c**). Same as in (**a**) for *MTOR* exon 45. (**d**) Same as in (**a**) for *ATP1A1* exon 4 (Q118R) from all single cell-derived K562 clones. Nucleotide substitutions are highlighted in bold. Insertions and deletions are highlighted in red, potential RT template and pegRNA scaffold incorporations in blue, and additional small locus duplications or uncharacterized insertions are highlighted in purple.

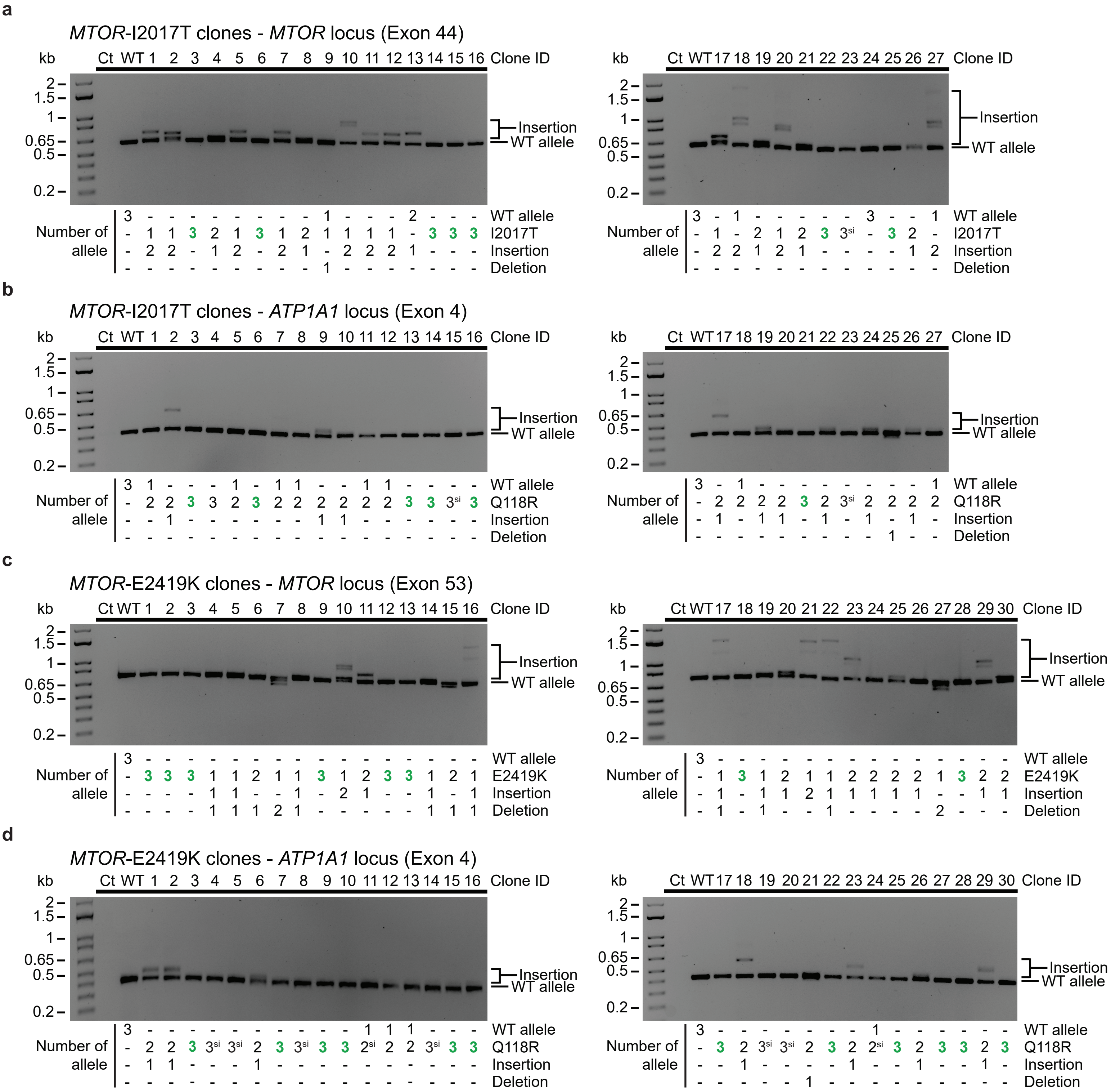
a
MTOR-I2017T clones - MTOR locus (Exon 44)
kb
2
1.5
1
0.65
0.5
0.2
Ct WT 1 2 3 4 5 6 7 8 9 10 11 12 13 14 15 16 Clone ID
Insertion
WT allele
Number of allele
3 - - - - - - - - 1 - - - 2 - - - WT allele
- 1 1 3 2 1 3 1 2 1 1 1 1 - 3 3 3 I2017T
- 2 2 - 1 2 - 2 1 - 2 2 2 1 - - - Insertion
- - - - - - - - - 1 - - - - - - - Deletion
Ct WT 17 18 19 20 21 22 23 24 25 26 27 Clone ID
3 - 1 - - - - - 3 - - 1 WT allele
- 1 - 2 1 2 3 $3^{si}$ - 3 2 - I2017T
- 2 2 1 2 1 - - - - 1 2 Insertion
- - - - - - - - - - - - Deletion
b
MTOR-I2017T clones - ATP1A1 locus (Exon 4)
Ct WT 1 2 3 4 5 6 7 8 9 10 11 12 13 14 15 16 Clone ID
3 1 - - - 1 - 1 1 - - 1 1 - - - - WT allele
- 2 2 3 3 2 3 2 2 2 2 2 2 3 3 $3^{si}$ 3 Q118R
- - 1 - - - - - - 1 1 - - - - - - Insertion
- - - - - - - - - - - - - - - - - Deletion
Ct WT 17 18 19 20 21 22 23 24 25 26 27 Clone ID
3 - 1 - - - - - - - - 1 WT allele
- 2 2 2 2 3 2 $3^{si}$ 2 2 2 2 Q118R
- 1 - 1 1 - 1 - 1 - 1 - Insertion
- - - - - - - - - 1 - - Deletion
c
MTOR-E2419K clones - MTOR locus (Exon 53)
Ct WT 1 2 3 4 5 6 7 8 9 10 11 12 13 14 15 16 Clone ID
3 - - - - - - - - - - - - - - - - WT allele
- 3 3 3 1 1 2 1 1 3 1 2 3 3 1 2 1 E2419K
- - - - 1 1 - - 1 - 2 1 - - 1 - 1 Insertion
- - - - 1 1 1 2 1 - - - - - 1 1 1 Deletion
Ct WT 17 18 19 20 21 22 23 24 25 26 27 28 29 30 Clone ID
3 - - - - - - - - - - - - - - WT allele
- 1 3 1 2 1 1 2 2 2 2 1 3 2 2 E2419K
- 1 - 1 1 2 1 1 1 1 1 - - 1 1 Insertion
- 1 - 1 - - 1 - - - - 2 - - - Deletion
d
MTOR-E2419K clones - ATP1A1 locus (Exon 4)
Ct WT 1 2 3 4 5 6 7 8 9 10 11 12 13 14 15 16 Clone ID
3 - - - - - - - - - - 1 1 1 - - - WT allele
- 2 2 3 $3^{si}$ $3^{si}$ 2 3 $3^{si}$ 3 3 $2^{si}$ 2 2 $3^{si}$ 3 3 Q118R
- 1 1 - - - 1 - - - - - - - - - - Insertion
- - - - - - - - - - - - - - - - - Deletion
Ct WT 17 18 19 20 21 22 23 24 25 26 27 28 29 30 Clone ID
3 - - - - - - - 1 - - - - - - WT allele
- 3 2 $3^{si}$ $3^{si}$ 2 3 2 $2^{si}$ 3 2 3 3 2 3 Q118R
- - 1 - - - - 1 - - 1 - - 1 - Insertion
- - - - - 1 - - - - - - - - - Deletion

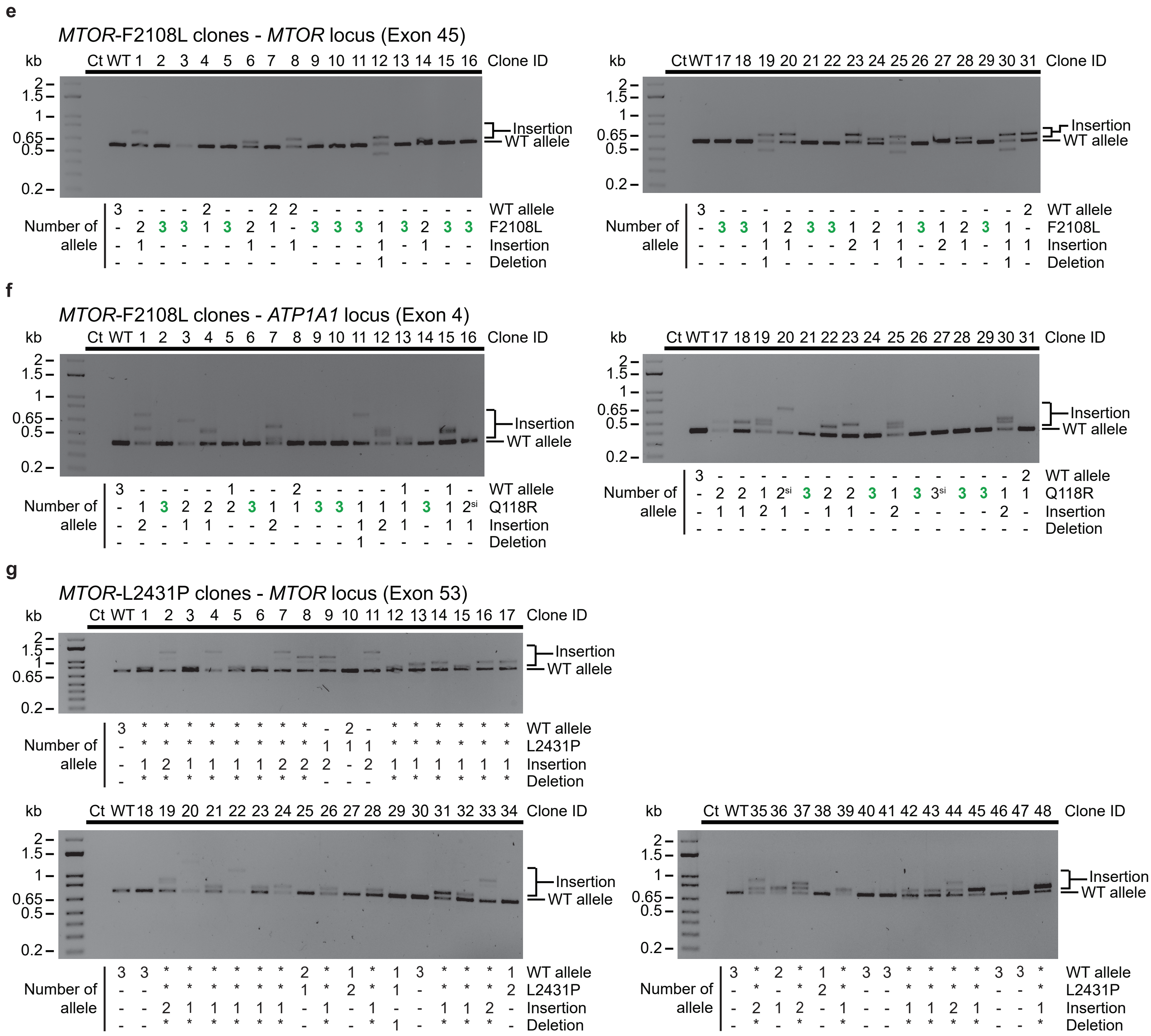
e
MTOR-F2108L clones - MTOR locus (Exon 45)
kb
2
1.5
1
0.65
0.5
0.2
Ct WT 1 2 3 4 5 6 7 8 9 10 11 12 13 14 15 16 Clone ID
Insertion
WT allele
Number of allele
3 - - - 2 - - 2 2 - - - - - - - - WT allele
- 2 3 3 1 3 2 1 - 3 3 3 1 3 2 3 3 F2108L
- 1 - - - - 1 - 1 - - - 1 - 1 - - Insertion
- - - - - - - - - - - 1 - - - - Deletion
Ct WT 17 18 19 20 21 22 23 24 25 26 27 28 29 30 31 Clone ID
3 - - - - - - - - - - - - - - 2 WT allele
- 3 3 1 2 3 3 1 2 1 3 1 2 3 1 - F2108L
- - - 1 1 - - 2 1 1 - 2 1 - 1 1 Insertion
- - - 1 - - - - - 1 - - - - 1 - Deletion
f
MTOR-F2108L clones - ATP1A1 locus (Exon 4)
Ct WT 1 2 3 4 5 6 7 8 9 10 11 12 13 14 15 16 Clone ID
3 - - - - 1 - - 2 - - - - 1 - 1 - WT allele
- 1 3 2 2 2 3 1 1 3 3 1 1 1 3 1 2si Q118R
- 2 - 1 1 - - 2 - - - 1 2 1 - 1 1 Insertion
- - - - - - - - - - - 1 - - - - - Deletion
Ct WT 17 18 19 20 21 22 23 24 25 26 27 28 29 30 31 Clone ID
3 - - - - - - - - - - - - - - 2 WT allele
- 2 2 1 2si 3 2 2 3 1 3 3si 3 3 1 1 Q118R
- 1 1 2 1 - 1 1 - 2 - - - - 2 - Insertion
- - - - - - - - - - - - - - - - Deletion
g
MTOR-L2431P clones - MTOR locus (Exon 53)
Ct WT 1 2 3 4 5 6 7 8 9 10 11 12 13 14 15 16 17 Clone ID
3 * * * * * * * * - 2 - * * * * * * WT allele
- * * * * * * * * 1 1 1 * * * * * * L2431P
- 1 2 1 1 1 1 2 2 2 - 2 1 1 1 1 1 1 Insertion
- * * * * * * * * - - - * * * * * * Deletion
Ct WT 18 19 20 21 22 23 24 25 26 27 28 29 30 31 32 33 34 Clone ID
3 3 * * * * * * 2 * 1 * 1 3 * * * 1 WT allele
- - * * * * * * 1 * 2 * 1 - * * * 2 L2431P
- - 2 1 1 1 1 1 - 1 - 1 - - 1 1 2 - Insertion
- - * * * * * * - * - * 1 - * * * - Deletion
Ct WT 35 36 37 38 39 40 41 42 43 44 45 46 47 48 Clone ID
3 * 2 * 1 * 3 3 * * * * 3 3 * WT allele
- * - * 2 * - - * * * * - - * L2431P
- 2 1 2 - 1 - - 1 1 2 1 - - 1 Insertion
- * - * - * - - * * * * - - * Deletion

**Supplementary figure 12 (Related to Fig. 3 and Table 1). Large PE3-mediated insertions in single cell-derived clones.** (**a**) PCR-based genotyping of *MTOR* exon 44 from single cell-derived K562 clones harboring the *MTOR*-I2017T hyperactivating mutation. (**b**) Same as in (**a**) for *ATP1A1* exon 4. (**c**) PCR-based genotyping of *MTOR* exon 53 from single cell-derived K562 clones harboring the *MTOR*-E2419K hyperactivating mutation. (**d**) Same as in (**c**) for *ATP1A1* exon 4. (**e**) PCR-based genotyping of *MTOR* exon 45 from single cell-derived K562 clones harboring the *MTOR*-F2108L rapamycin resistance mutation. (**f**) Same as in (**e**) for *ATP1A1* exon 4. (**g**) PCR-based genotyping of *MTOR* exon 53 from single cell-derived K562 clones harboring the *MTOR*-L2431P hyperactivating mutation. The number of precise prime edited alleles was determined from BEAT Sanger sequencing trace analysis and small insertions and deletions were analysed with DECODR. Larger insertions are indicated, and homozygous single cell-derived clones are highlighted in bold and green. PE alleles harboring pegRNA scaffold incorporation are indicated (si). PCR products that were not sent for sequencing (*MTOR*-L2431P clones) are highlighted with a star.

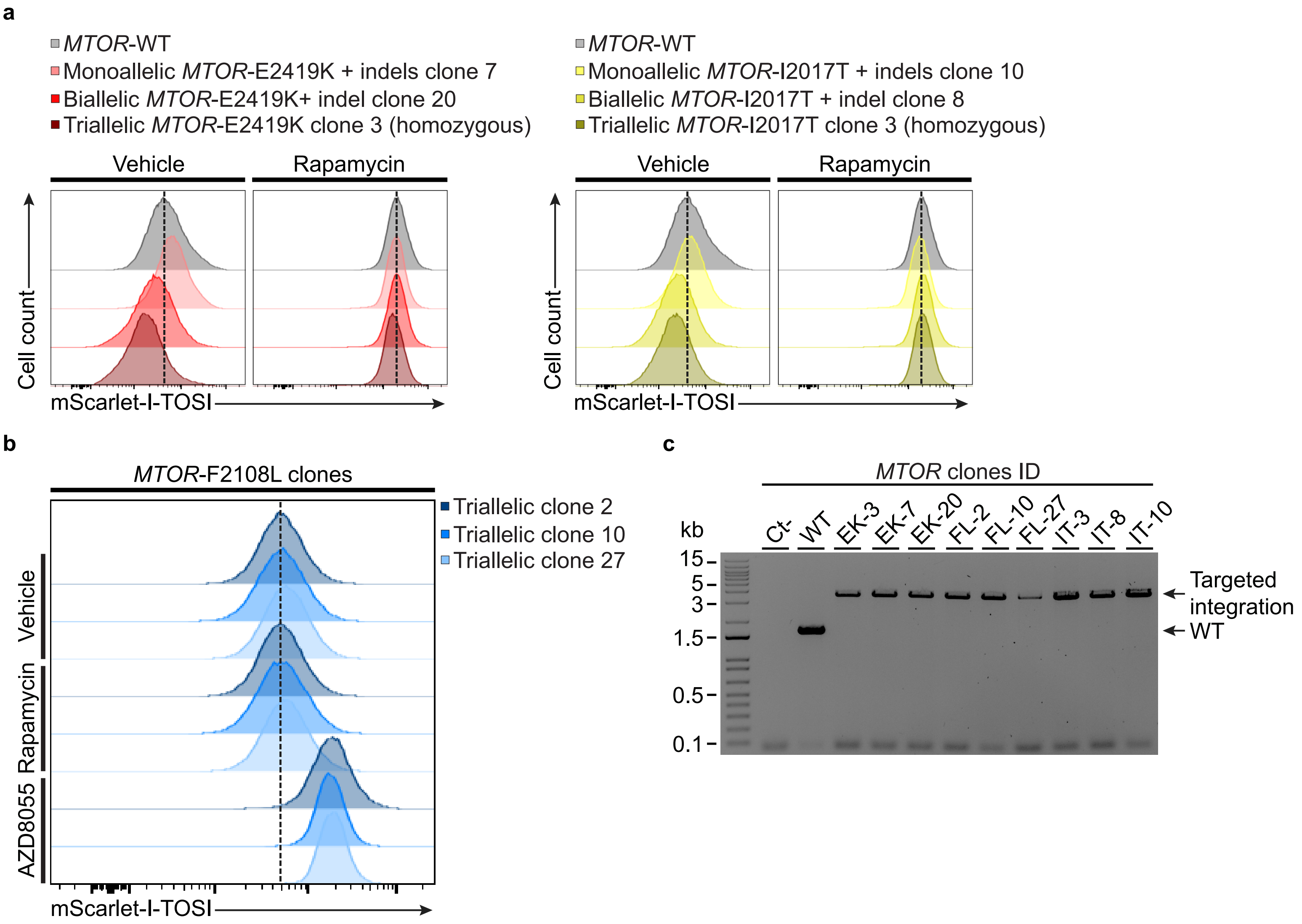
a
MTOR-WT
Monoallelic MTOR-E2419K + indels clone 7
Biallelic MTOR-E2419K+ indel clone 20
Triallelic MTOR-E2419K clone 3 (homozygous)
MTOR-WT
Monoallelic MTOR-I2017T + indels clone 10
Biallelic MTOR-I2017T + indel clone 8
Triallelic MTOR-I2017T clone 3 (homozygous)
Vehicle
Rapamycin
Cell count
mScarlet-I-TOSI
b
MTOR-F2108L clones
Triallelic clone 2
Triallelic clone 10
Triallelic clone 27
Vehicle
Rapamycin
AZD8055
mScarlet-I-TOSI
c
MTOR clones ID
kb
Ct-
WT
EK-3
EK-7
EK-20
FL-2
FL-10
FL-27
IT-3
IT-8
IT-10
15
5
3
1.5
0.5
0.1
Targeted integration
WT

**Supplementary figure 13 (Related to Fig. 3 and Table 1). mSc-TOSI reporter intensity in single cell-derived clones harboring *MTOR* mutations.** (**a**) Histogram plot of mSc-TOSI intensity in monoallelic, biallelic, and triallelic (homozygous) single cell-derived K562 clones harboring *MTOR* hyperactivating mutations. (**b**) Same as in (**a**) with homozygous clones harboring the *MTOR*-F2108L rapamycin resistance mutation. After coselection, cells were treated for 24 hours with 50 nM rapamycin or 50 nM AZD8055 before FACS analysis. Representative FACS images are from one of two independent biological replicates performed at different times with equivalent results (See **Fig. 3**). (**c**) Out-Out PCR for mSc-TOSI knock-in detection at *ATP1A1* intron 17 in single cell-derived clones.

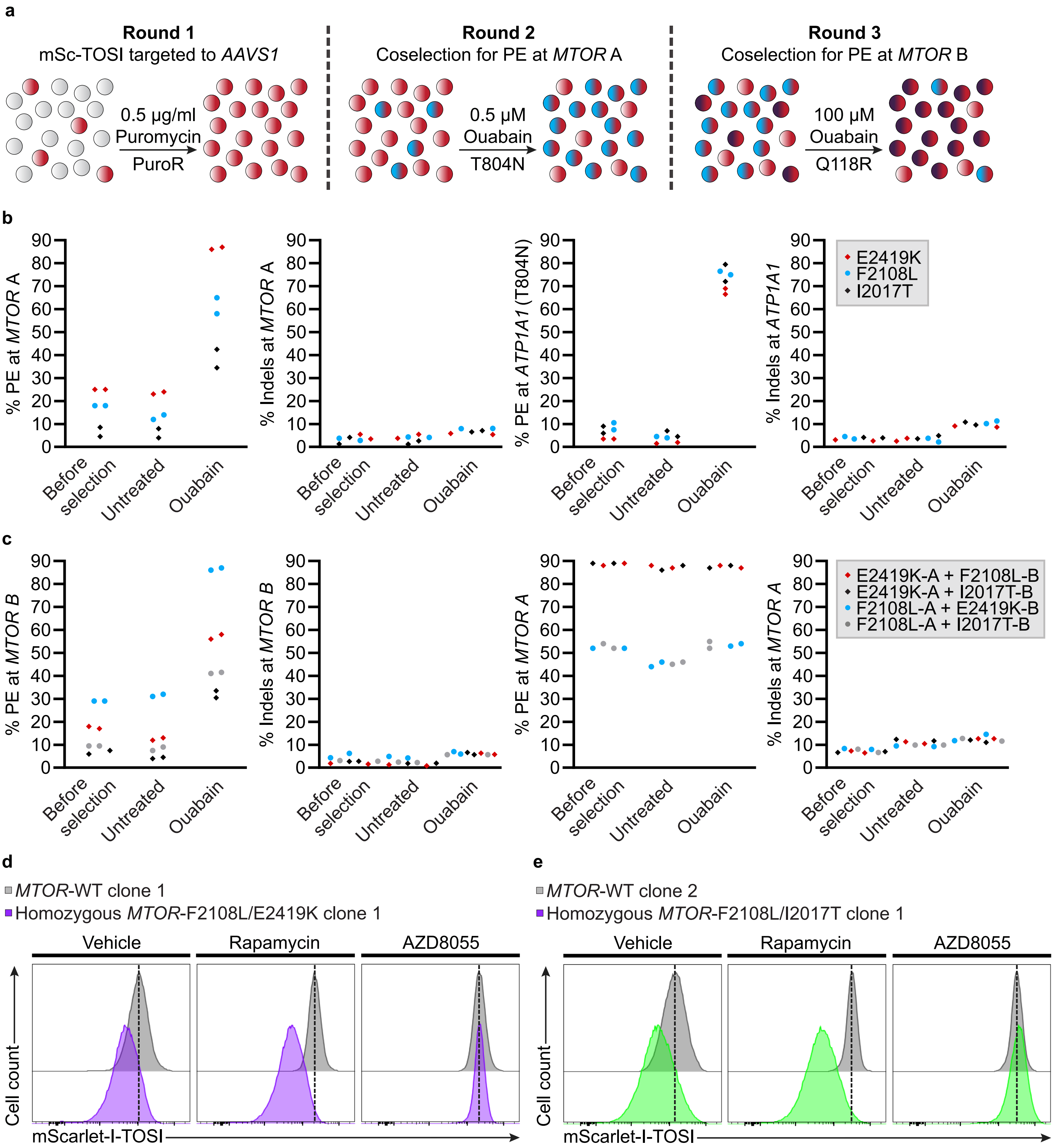
a
Round 1
mSc-TOSI targeted to AAVS1
0.5 µg/ml
Puromycin
PuroR
Round 2
Coselection for PE at MTOR A
0.5 µM
Ouabain
T804N
Round 3
Coselection for PE at MTOR B
100 µM
Ouabain
Q118R
b
% PE at MTOR A
% Indels at MTOR A
% PE at ATP1A1 (T804N)
% Indels at ATP1A1
E2419K
F2108L
I2017T
Before selection
Untreated
Ouabain
c
% PE at MTOR B
% Indels at MTOR B
% PE at MTOR A
% Indels at MTOR A
E2419K-A + F2108L-B
E2419K-A + I2017T-B
F2108L-A + E2419K-B
F2108L-A + I2017T-B
d
MTOR-WT clone 1
Homozygous MTOR-F2108L/E2419K clone 1
Vehicle
Rapamycin
AZD8055
Cell count
mScarlet-I-TOSI
e
MTOR-WT clone 2
Homozygous MTOR-F2108L/I2017T clone 1
Vehicle
Rapamycin
AZD8055
Cell count
mScarlet-I-TOSI

**Supplementary figure 14. Successive rounds of coselection for the isolation of double *MTOR* mutants.** (**a**) Schematic representation of reporter cell line engineering. K562 cells stably expressing the mSc-TOSI reporter from the *AAVS1* locus are first selected with 0.5 µg/ml puromycin (see **Supplementary Fig. 9**). Following the first round of selection with puromycin, coselection for PE to install the first mutation at *MTOR* (*MTOR* A) is performed with 0.5 µM ouabain. Then, a successive round of coselection for PE to install the second mutation at *MTOR* (*MTOR* B) is performed with 100 µM ouabain. (**b**) PE and small indels quantification as determined by BEAT and TIDE analysis from Sanger sequencing. K562 cells stably expressing the mSc-TOSI reporter from the *AAVS1* locus were transfected with PE3 vectors targeting *ATP1A1* exon 17 (T804N) and *MTOR* A. Genomic DNA was harvested 3 days post-transfection (before selection) and cells were treated (ouabain) or not (untreated) with 0.5 µM ouabain until all non-resistant cells were eliminated. $n$ = 2 independent biological replicates performed at different times. (**c**) Same as in (**b**) for a successive round of coselection for PE at *MTOR* B with K562 cells harboring the *ATP1A1*-T804N and the indicated *MTOR* A mutation. Successive coselection was performed with 100 µM ouabain until all non-resistant cells were eliminated. $n$ = 2 independent biological replicates performed at different times. (**d**) Histogram plot of mSc-TOSI intensity in a homozygous single cell-derived K562 clone harboring the *MTOR*-F2108L rapamycin resistance mutation and the *MTOR*-E2419K hyperactivating mutation. (**e**) Same as in (**d**) with a homozygous clone harboring the *MTOR*-F2108L rapamycin resistance mutation and the *MTOR*-I2017T hyperactivating mutation. Where indicated, cells were treated for 24 hours with 50 nM rapamycin or 50 nM AZD8055 before FACS analysis. Representative images are from one of two independent biological replicates performed at different times with equivalent results (see **Fig. 3**).

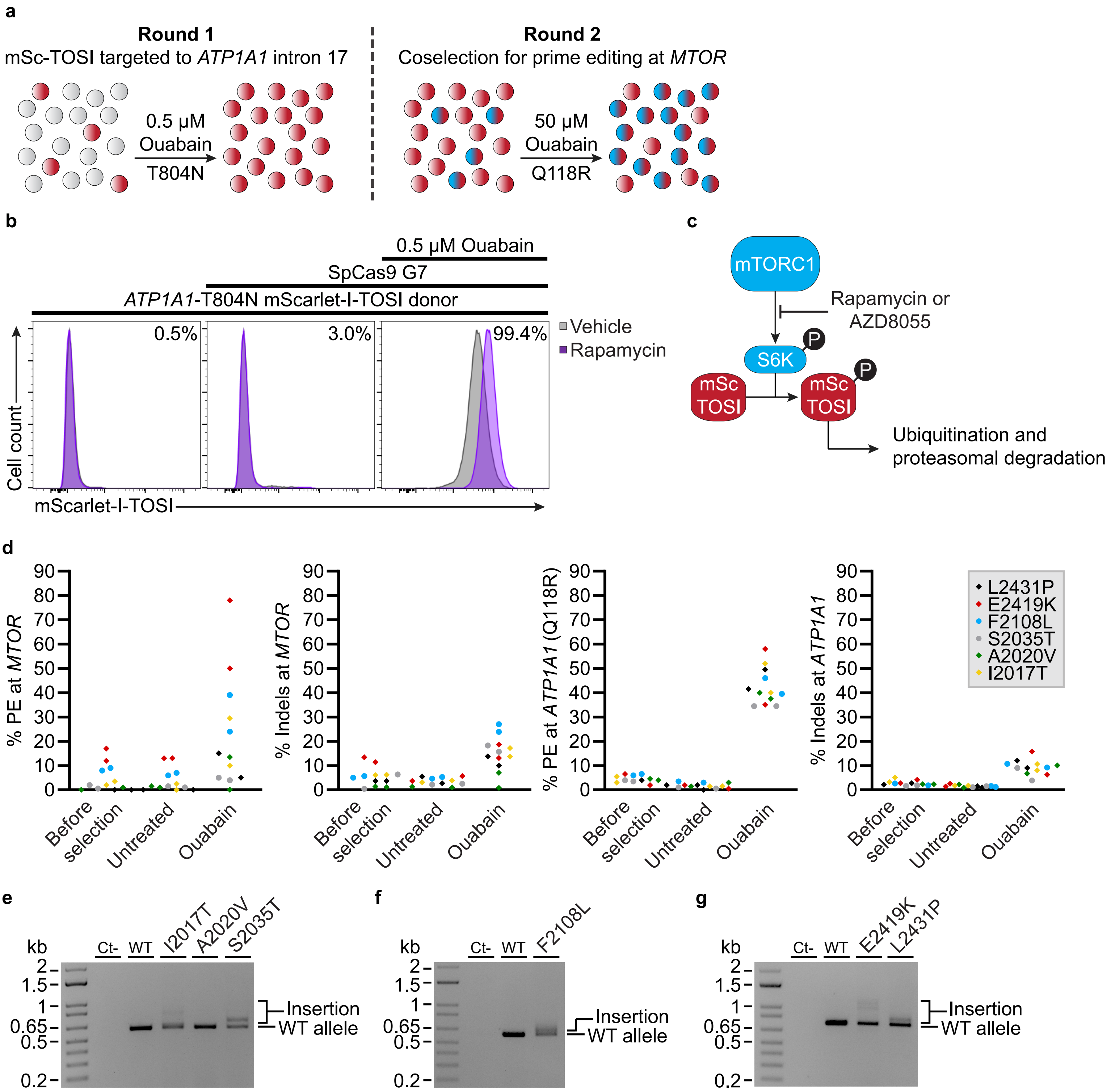

a
Round 1
mSc-TOSI targeted to ATP1A1 intron 17
0.5 μM Ouabain
T804N
Round 2
Coselection for prime editing at MTOR
50 μM Ouabain
Q118R
b
0.5 μM Ouabain
SpCas9 G7
ATP1A1-T804N mScarlet-I-TOSI donor
0.5%
3.0%
99.4%
Vehicle
Rapamycin
Cell count
mScarlet-I-TOSI
c
mTORC1
Rapamycin or AZD8055
S6K
P
mSc TOSI
Ubiquitination and proteasomal degradation
d
% PE at MTOR
% Indels at MTOR
% PE at ATP1A1 (Q118R)
% Indels at ATP1A1
Before selection
Untreated
Ouabain
L2431P
E2419K
F2108L
S2035T
A2020V
I2017T
e
kb
Ct-
WT
I2017T
A2020V
S2035T
Insertion
WT allele
f
F2108L
g
E2419K
L2431P

**Supplementary figure 15 (Related to Fig. 3). Sequential mSc-TOSI knock-in and coselection for prime editing at *MTOR* in HeLa S3 cells.** (**a**) Schematic representation of reporter cell line engineering. HeLa S3 cells stably expressing the mSc-TOSI reporter from *ATP1A1* intron 17 are first selected with 0.5 µM ouabain. Then, coselection for PE to insert the indicated mutations at *MTOR* is performed with 50 µM ouabain. (**b**) FACS-based quantification of mSc-TOSI knock-in at *ATP1A1* intron 17. HeLa S3 cells were transfected with SpCas9 and a donor targeting *ATP1A1* and treated with or without 0.5 µM ouabain 3 days post-transfection until all non-resistant cells were eliminated. Where indicated, cells were treated for 24 hours with 500 nM rapamycin before FACS analysis. Representative FACS images are from one of two independent biological replicates performed at different times with equivalent results. (**c**) Schematic of mSc-TOSI degradation under mTORC1 signaling. In these cells the difference in fluorescence observed by FACS-based analysis was too minimal between the on and off states (+/- rapamycin) to monitor the impact of PE-induced mutations. (**d**) PE and small indels quantification as determined by BEAT and TIDE analysis from Sanger sequences. HeLa S3 cells stably expressing the mSc-TOSI reporter were transfected with PE3 vectors targeting *ATP1A1* exon 4 (Q118R) and *MTOR*. Genomic DNA was harvested 3 days post-transfection (before selection) and cells were treated (ouabain) or not (untreated) with 50 µM ouabain until all non-resistant cells were eliminated. $n$ = 2 independent biological replicates performed at different times. (**e**) PCR-based detection of large PE-mediated insertions at *MTOR* exon 44 in bulk coselected populations of HeLa S3 cells. (**f**) Same as in (**e**) at *MTOR* exon 45. (**g**) Same as in (**e**) at *MTOR* exon 53.

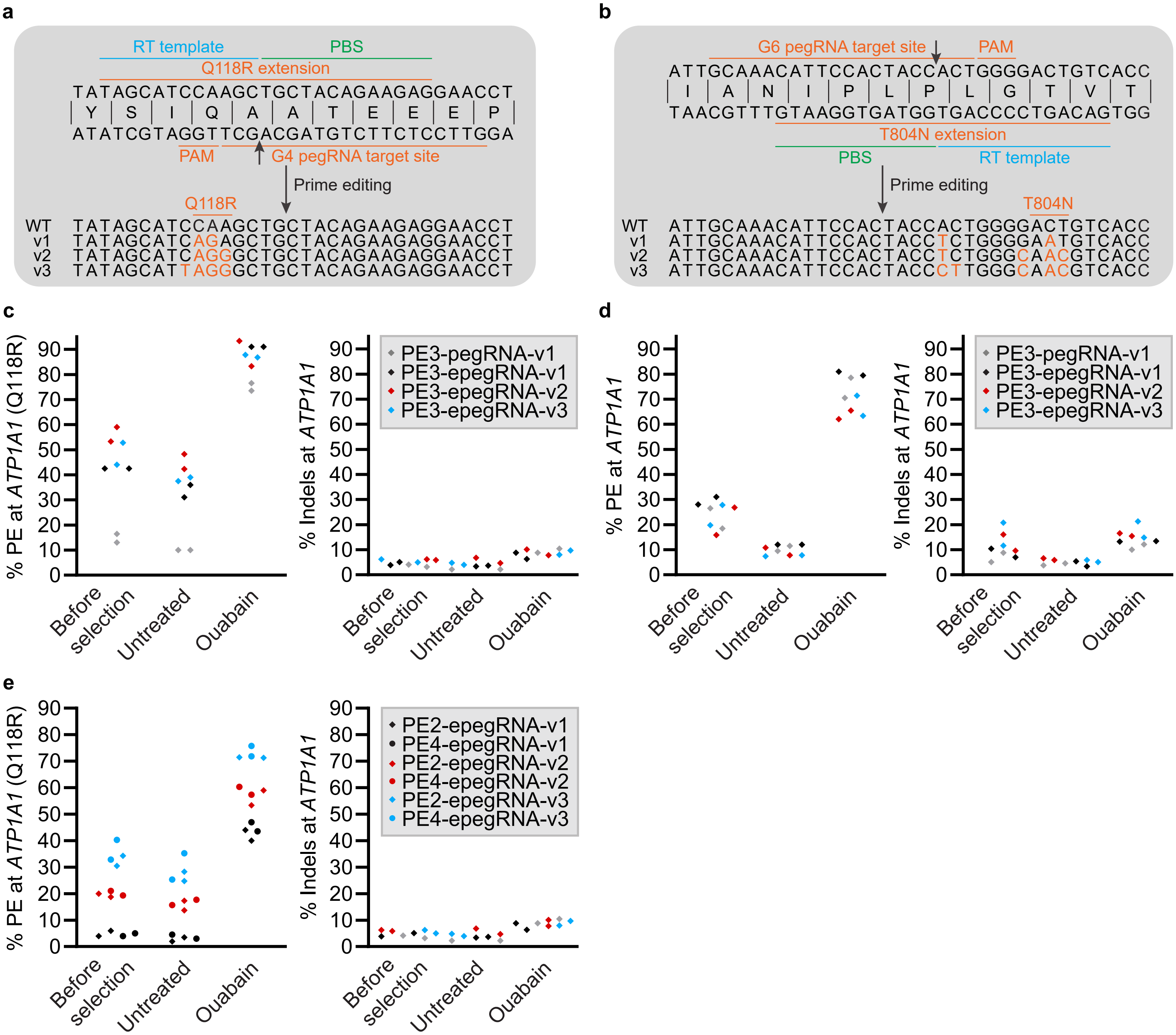

a
RT template
PBS
Q118R extension
TATAGCATCCAAGCTGCTACAGAAGAGGAACCT
Y S I Q A A T E E E P
ATATCGTAGGTTCGACGATGTCTTCTCCTTGGA
PAM
G4 pegRNA target site
Prime editing
Q118R
WT TATAGCATCCAAGCTGCTACAGAAGAGGAACCT
v1 TATAGCATCAGAGCTGCTACAGAAGAGGAACCT
v2 TATAGCATCAGGGCTGCTACAGAAGAGGAACCT
v3 TATAGCATTAGGGCTGCTACAGAAGAGGAACCT
b
G6 pegRNA target site
PAM
ATTGCAAACATTCCACTACCACTGGGGACTGTCACC
I A N I P L P L G T V T
TAACGTTTGTAAGGTGATGGTGACCCCTGACAGTGG
T804N extension
PBS
RT template
Prime editing
T804N
WT ATTGCAAACATTCCACTACCACTGGGGACTGTCACC
v1 ATTGCAAACATTCCACTACCTCTGGGGAATGTCACC
v2 ATTGCAAACATTCCACTACCTCTGGGCAACGTCACC
v3 ATTGCAAACATTCCACTACCCTTGGGCAACGTCACC
c
% PE at ATP1A1 (Q118R)
% Indels at ATP1A1
PE3-pegRNA-v1
PE3-epegRNA-v1
PE3-epegRNA-v2
PE3-epegRNA-v3
Before selection
Untreated
Ouabain
d
% PE at ATP1A1
% Indels at ATP1A1
PE3-pegRNA-v1
PE3-epegRNA-v1
PE3-epegRNA-v2
PE3-epegRNA-v3
Before selection
Untreated
Ouabain
e
% PE at ATP1A1 (Q118R)
% Indels at ATP1A1
PE2-epegRNA-v1
PE4-epegRNA-v1
PE2-epegRNA-v2
PE4-epegRNA-v2
PE2-epegRNA-v3
PE4-epegRNA-v3
Before selection
Untreated
Ouabain
0
10
20
30
40
50
60
70
80
90

**Supplementary figure 16. Engineered pegRNAs (epegRNAs) with MMR-evading mutations enhance prime editing at *ATP1A1*.** (**a**) Schematic representation of the SpCas9 target site and the reverse transcriptase (RT) template used to install the Q118R mutation at *ATP1A1* exon 4 to induce cellular resistance to ouabain. New epegRNA versions harboring additional MMR-evading mutations are shown. (**b**) Schematic representation of the SpCas9 target site and the reverse transcriptase (RT) template used to install the T804N mutation at *ATP1A1* exon 17 to induce cellular resistance to ouabain. New epegRNA versions harboring additional MMR-evading mutations are shown. (**c**) PE and small indels quantification at *ATP1A1* exon 4 (Q118R) as determined by BEAT and TIDE analysis from Sanger sequences. K562 cells were transfected with PE3max vectors and genomic DNA was harvested 3 days post-transfection (before selection) and cells were treated (ouabain) or not (untreated) with 0.5 µM ouabain for 10 days starting 3 days post-transfection. $n = 2$ independent biological replicates performed at different times. (**d**) Same as in (**c**) at *ATP1A1* exon 17 (T804N). (**e**) Same as in (**c**), but K562 cells were transfected with PEmax (PE2) or PEmax-2A-hMLH1dn (PE4) and pegRNA vectors without an additional nick sgRNA.

a

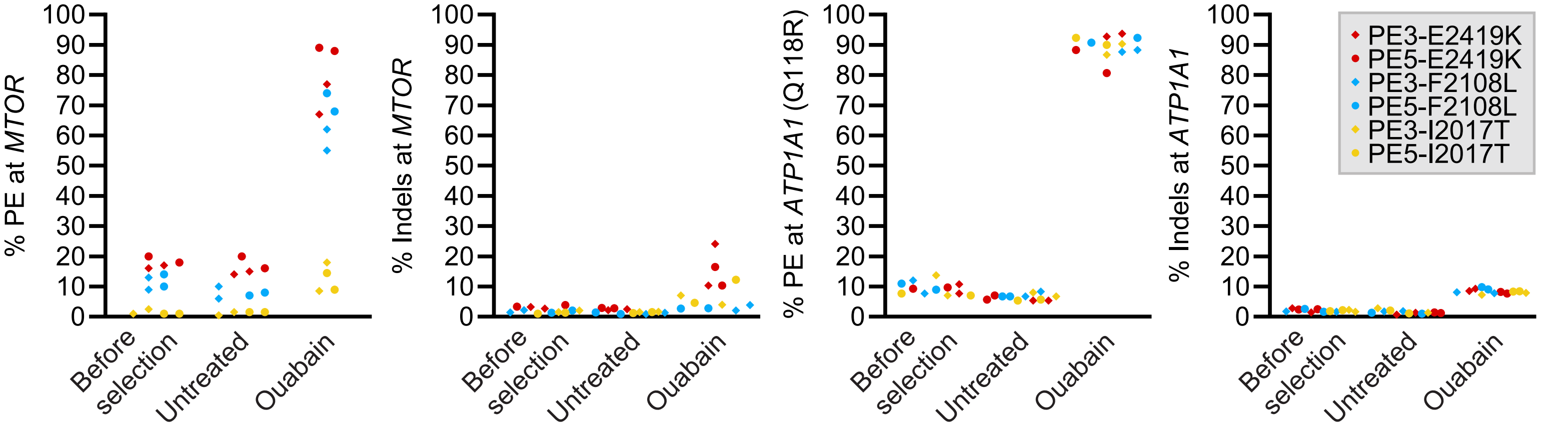

% PE at MTOR
% Indels at MTOR
% PE at ATP1A1 (Q118R)
% Indels at ATP1A1
0
10
20
30
40
50
60
70
80
90
100
Before selection
Untreated
Ouabain
PE3-E2419K
PE5-E2419K
PE3-F2108L
PE5-F2108L
PE3-I2017T
PE5-I2017T

**Supplementary figure 17 (Related to Fig. 3). Robust coselection for prime editing at *MTOR* in U2OS cells with engineered pegRNAs (epegRNAs).** (**a**) PE and small indels quantification as determined by BEAT and TIDE analysis from Sanger sequences. U2OS cells were transfected with PE3max-epegRNA vectors targeting *ATP1A1* exon 4 (epegRNA-Q118R_v2) and *MTOR*. Genomic DNA was harvested 3 days post-transfection (before selection) and cells were treated (ouabain) or not (untreated) with 0.5 µM ouabain until all non-resistant cells were eliminated. $n$ = 2 independent biological replicates performed at different times.

a

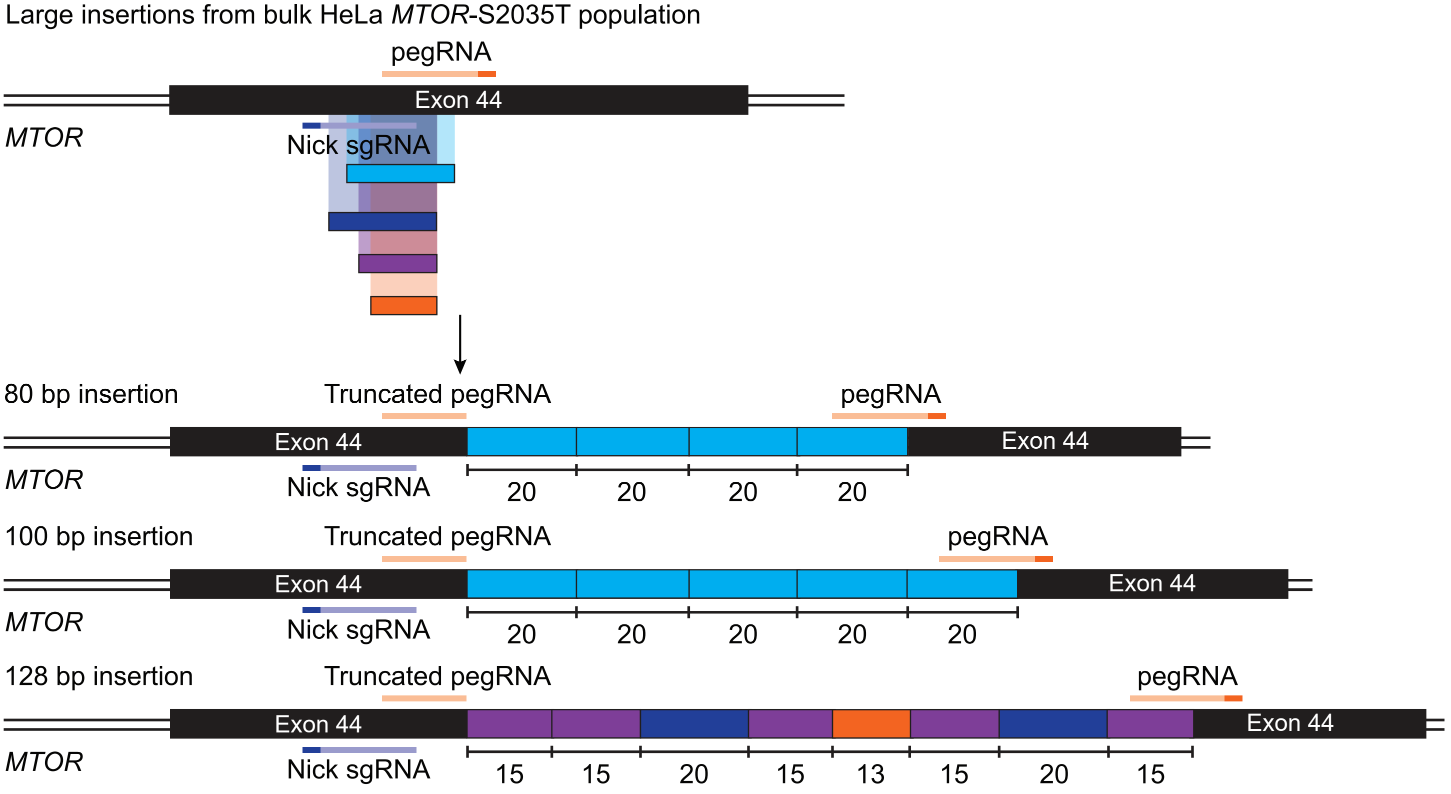

Large insertions from bulk HeLa *MTOR*-S2035T population
pegRNA
Exon 44
*MTOR*
Nick sgRNA
80 bp insertion
Truncated pegRNA
pegRNA
Exon 44
Exon 44
*MTOR*
Nick sgRNA
20 20 20 20
100 bp insertion
Truncated pegRNA
pegRNA
Exon 44
Exon 44
*MTOR*
Nick sgRNA
20 20 20 20 20
128 bp insertion
Truncated pegRNA
pegRNA
Exon 44
Exon 44
*MTOR*
Nick sgRNA
15 15 20 15 13 15 20 15


b

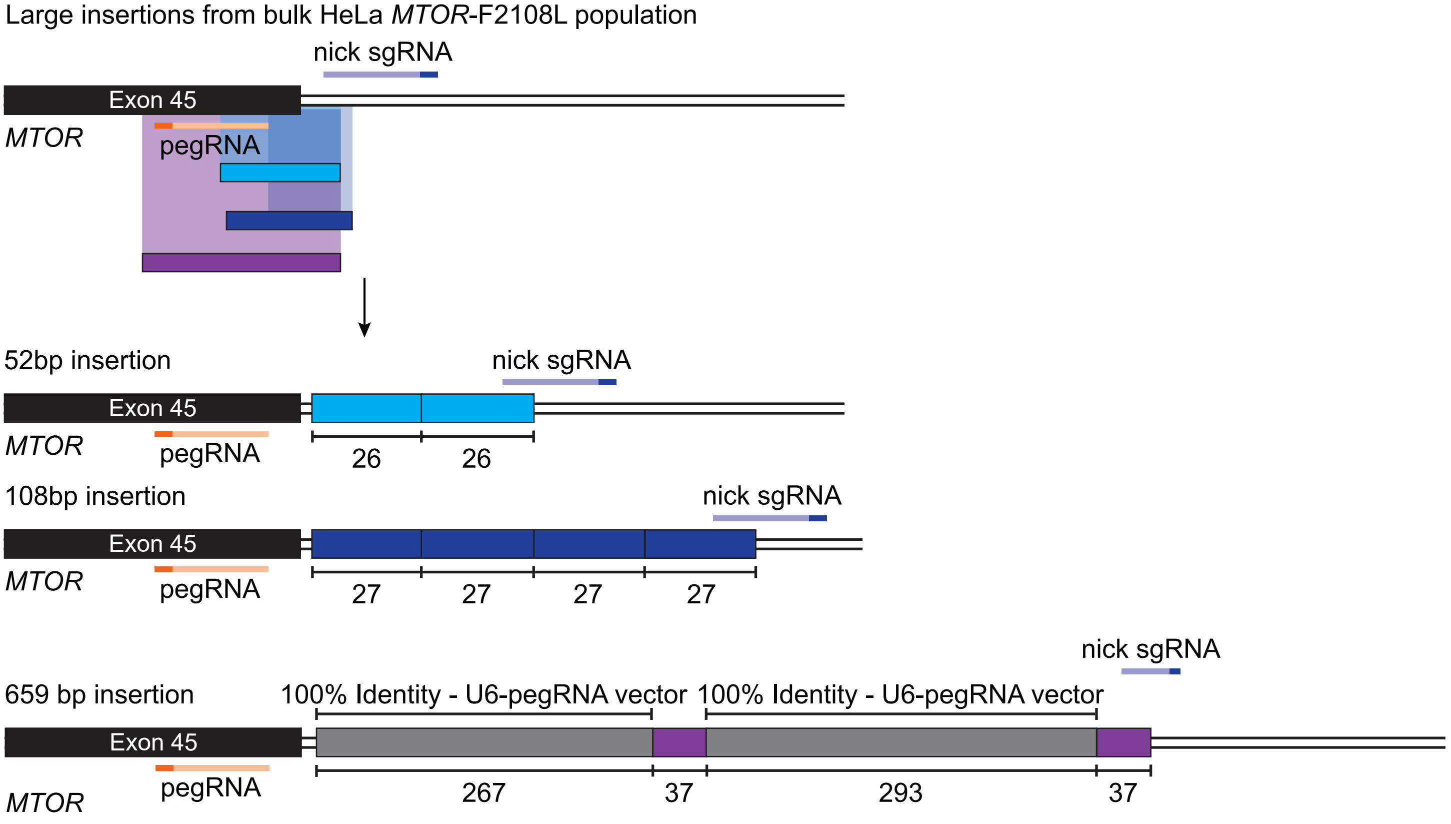

Large insertions from bulk HeLa *MTOR*-F2108L population
nick sgRNA
Exon 45
*MTOR*
pegRNA
52bp insertion
nick sgRNA
Exon 45
*MTOR*
pegRNA
26 26
108bp insertion
nick sgRNA
Exon 45
*MTOR*
pegRNA
27 27 27 27
nick sgRNA
659 bp insertion
100% Identity - U6-pegRNA vector
100% Identity - U6-pegRNA vector
Exon 45
pegRNA
*MTOR*
267 37 293 37

**Supplementary figure 18 (Related to Supplementary Fig. 15). Characterization of large PE3-mediated insertions in bulk populations of HeLa S3 cells.** (**a**) Schematic representation of the large insertions observed in HeLa S3 cells following coselection for PE at *MTOR* exon 44 for the installation of the *MTOR*-S2035T mutation. As described in **Supplementary Fig. 15**, HeLa S3 cells stably expressing the mSc-TOSI reporter were transfected with PE3 vectors targeting *ATP1A1* exon 4 (Q118R) and *MTOR*. Genomic DNA was harvested after ouabain coselection. TOPO cloning and Sanger sequencing were performed to characterize the large insertion. (**b**) Same as in (**a**) following coselection for PE at *MTOR* exon 45 for the installation of the *MTOR*-F2108L mutation.

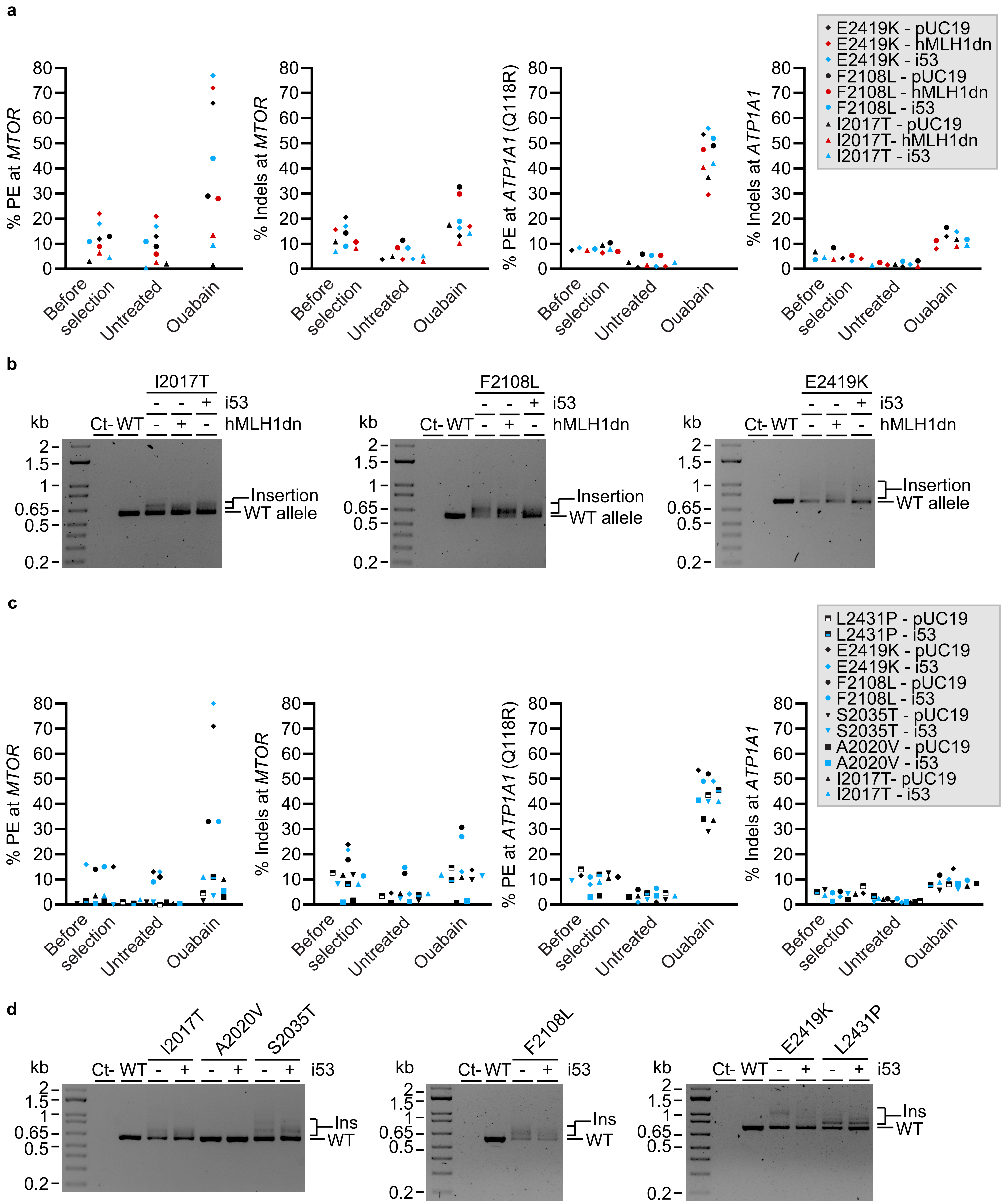
a
E2419K - pUC19
E2419K - hMLH1dn
E2419K - i53
F2108L - pUC19
F2108L - hMLH1dn
F2108L - i53
I2017T - pUC19
I2017T- hMLH1dn
I2017T - i53
% PE at MTOR
% Indels at MTOR
% PE at ATP1A1 (Q118R)
% Indels at ATP1A1
0
10
20
30
40
50
60
70
80
Before selection
Untreated
Ouabain
b
I2017T
F2108L
E2419K
i53
hMLH1dn
kb
Ct-
WT
2
1.5
1
0.65
0.5
0.2
Insertion
WT allele
c
L2431P - pUC19
L2431P - i53
E2419K - pUC19
E2419K - i53
F2108L - pUC19
F2108L - i53
S2035T - pUC19
S2035T - i53
A2020V - pUC19
A2020V - i53
I2017T- pUC19
I2017T - i53
d
I2017T
A2020V
S2035T
F2108L
E2419K
L2431P
Ins
WT

e

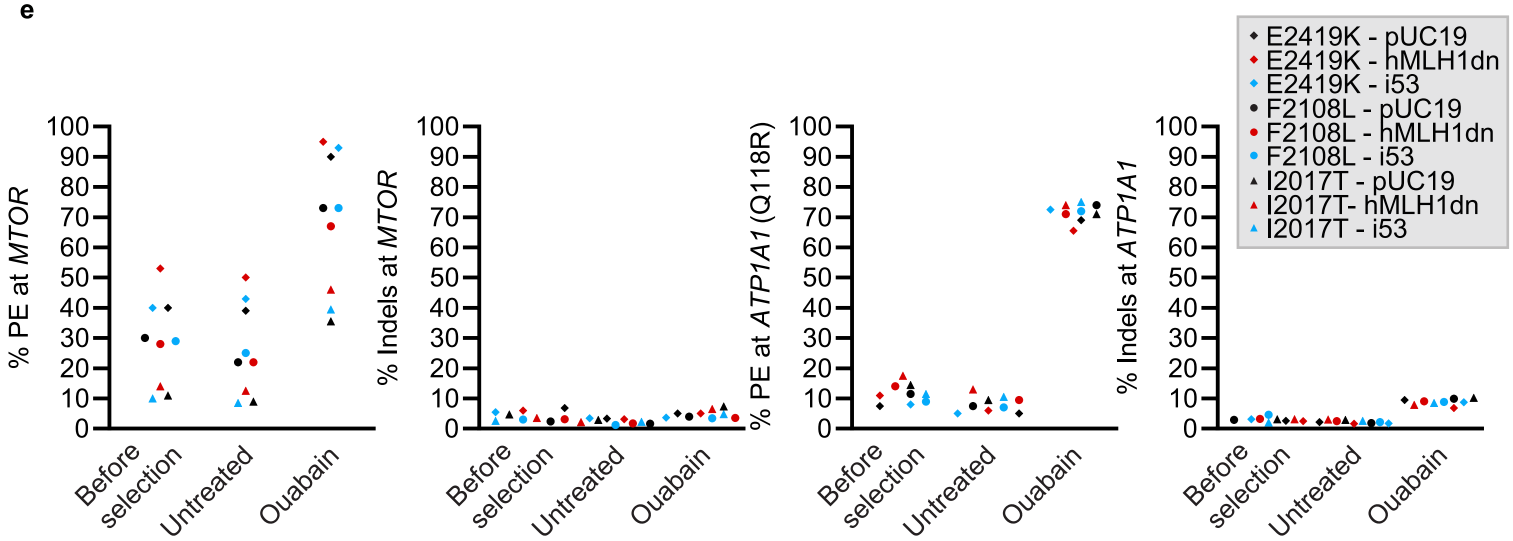

% PE at MTOR
% Indels at MTOR
% PE at ATP1A1 (Q118R)
% Indels at ATP1A1
0 10 20 30 40 50 60 70 80 90 100
Before selection
Untreated
Ouabain
E2419K - pUC19
E2419K - hMLH1dn
E2419K - i53
F2108L - pUC19
F2108L - hMLH1dn
F2108L - i53
I2017T - pUC19
I2017T- hMLH1dn
I2017T - i53

**Supplementary figure 19. Assessing the impact of 53BP1 inhibition on large PE-mediated insertions.** (**a**) PE and small indels quantification as determined by BEAT and TIDE analysis from Sanger sequences. HeLa S3 cells stably expressing the mSc-TOSI reporter were transfected with PE3max vectors targeting *ATP1A1* exon 4 (Q118R) and *MTOR*, and a vector expressing either hMLH1dn or i53. An empty pUC19 vector was used as a control to normalize DNA concentration in all transfections. Genomic DNA was harvested 3 days post-transfection (before selection) and cells were treated (ouabain) or not (untreated) with 50 µM ouabain until all non-resistant cells were eliminated. (**b**) PCR-based detection of large PE-mediated insertions at *MTOR* exon 44, 45, and 53 in bulk coselected populations of HeLa S3 cells. *n* = 1 experiment. (**c**) Experiment replicated as described in (**a**) using additional pegRNA/sgRNA pairs. (**d**) Same as in (**b**) with bulk populations of HeLa S3 cells from (**c**). *n* = 1 experiment. (**e**) PE and small indels quantification as determined by BEAT and TIDE analysis from Sanger sequences. K562 cells were transfected with PE3max vectors targeting *ATP1A1* exon 4 (Q118R) and *MTOR*, and a vector expressing either hMLH1dn or i53. An empty pUC19 vector was used as a control to normalize DNA concentration in all transfections. Genomic DNA was harvested 3 days post-transfection (before selection) and cells were treated (ouabain) or not (untreated) with 0.5 µM ouabain until all non-resistant cells were eliminated. *n* = 1 experiment.

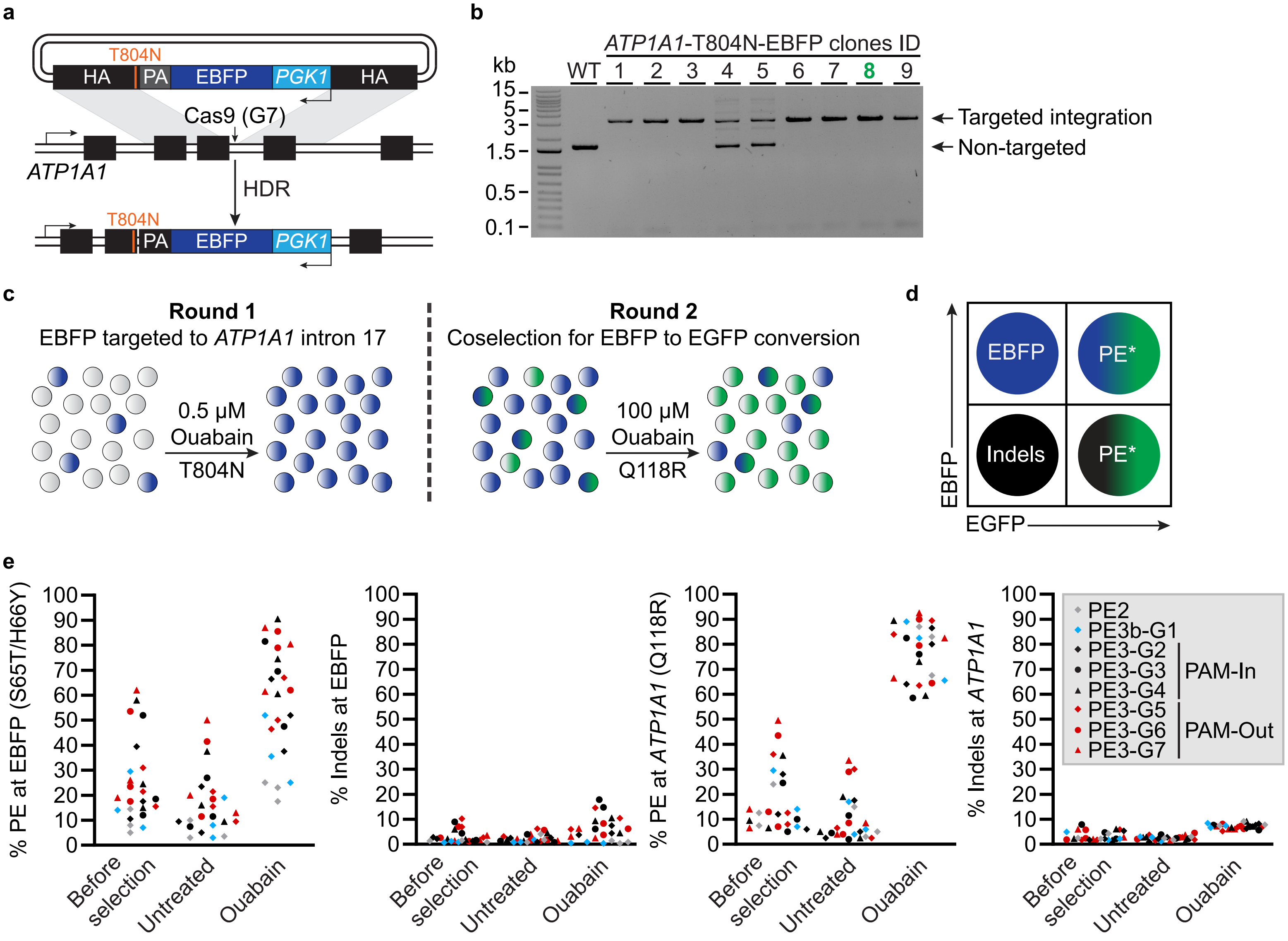
a
T804N
HA
PA
EBFP
PGK1
HA
Cas9 (G7)
ATP1A1
HDR
T804N
PA
EBFP
PGK1
b
ATP1A1-T804N-EBFP clones ID
kb
WT
1
2
3
4
5
6
7
8
9
15
5
3
1.5
0.5
0.1
Targeted integration
Non-targeted
c
Round 1
EBFP targeted to ATP1A1 intron 17
0.5 µM
Ouabain
T804N
Round 2
Coselection for EBFP to EGFP conversion
100 µM
Ouabain
Q118R
d
EBFP
PE*
Indels
PE*
EBFP
EGFP
e
% PE at EBFP (S65T/H66Y)
% Indels at EBFP
% PE at ATP1A1 (Q118R)
% Indels at ATP1A1
Before selection
Untreated
Ouabain
PE2
PE3b-G1
PE3-G2
PE3-G3
PE3-G4
PE3-G5
PE3-G6
PE3-G7
PAM-In
PAM-Out

**Supplementary figure 20 (Related to Fig. 6). Assessing the impact of complementary-strand nick locations on prime editing outcomes.** (**a**) Schematic of targeted EBFP integration at *ATP1A1* intron 17. (**b**) Out-Out PCR for EBFP knock-in detection at *ATP1A1* intron 17 in single cell-derived K562 clones. The *ATP1A1*-T804N-EBFP clone that was used during this study (clone 8) is highlighted in bold and green. (**c**) Schematic representation of reporter cell line engineering. K562 cells stably expressing the EBFP reporter from *ATP1A1* intron 17 are first selected with 0.5 µM ouabain. Then, coselection for EBFP to EGFP conversion via PE is performed with 100 µM ouabain. (**d**) Schematic of the FACS-based quantification of EBFP to EGFP conversion via PE. In K562 cells homozygous for EBFP integration at *ATP1A1* (triallelic), FACS-based analysis reports PE outcomes indirectly; (i) EBFP(+)/EGFP(+) cells result from monoallelic or biallelic PE, (ii) EBFP(-)/EGFP(+) cells are generated by either triallelic PE or combinations of mono- and biallelic PE along with indels*, (iii) EBFP(-)/EGFP(-) occur from indels on the three copies of the reporter. Indels are defined broadly in this context as any edits that inactivates the reporter. (**e**) PE and small indels quantification as determined by BEAT and TIDE analysis from Sanger sequencing. K562 cells stably expressing the EBFP reporter were transfected with PE3max vectors targeting *ATP1A1* exon 4 (pegRNA-Q118R_v1) and EBFP (epegRNA). Genomic DNA was harvested 3 days post-transfection (before selection) and cells were treated (ouabain) or not (untreated) with 100 µM ouabain until all non-resistant cells were eliminated. *n* = 3 independent biological replicates performed at different times. *PGK1*, human phosphoglycerate kinase 1 promoter. PA, polyadenylation signal. HA, homology arm.

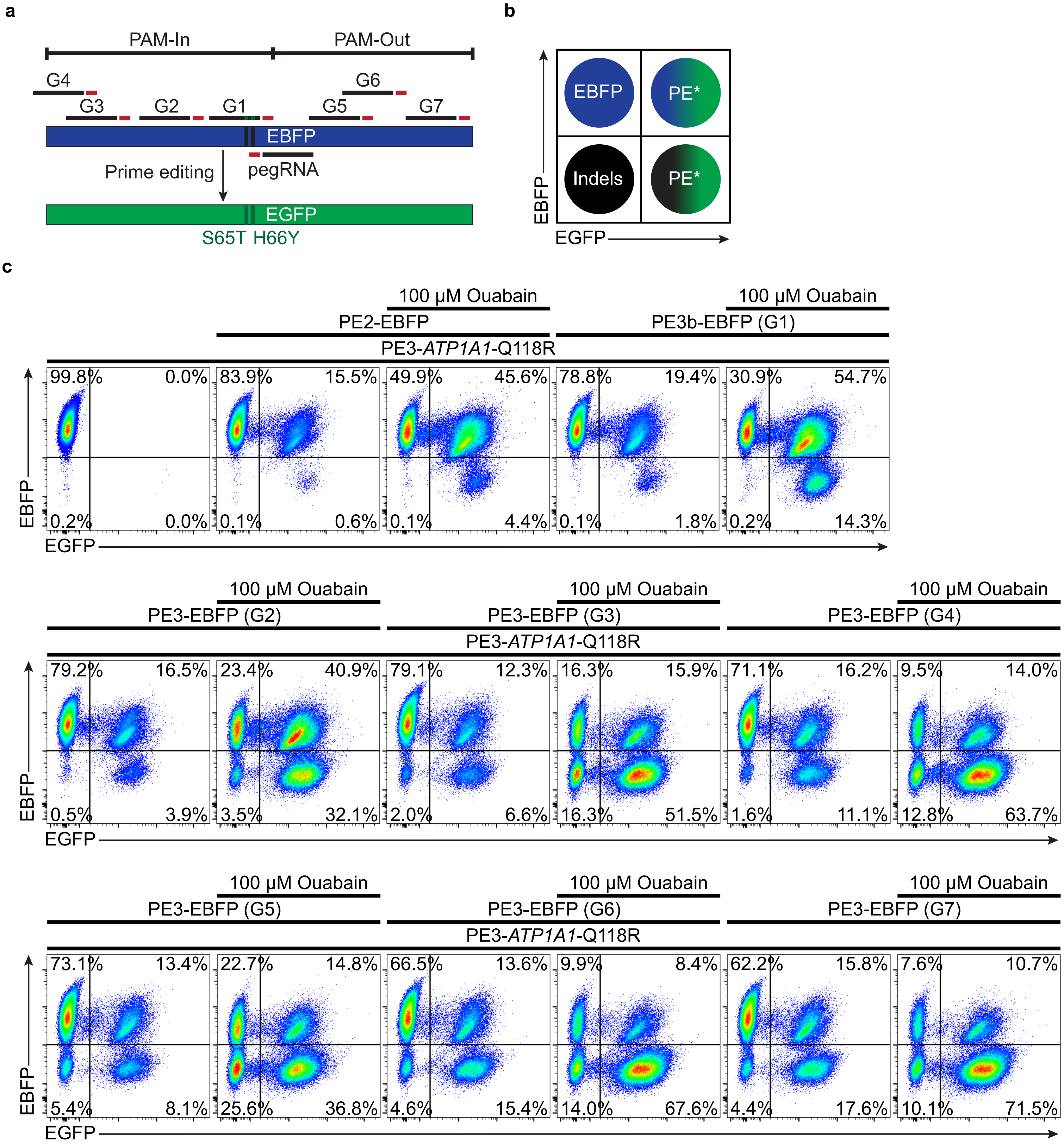
a
PAM-In
PAM-Out
G4
G6
G3
G2
G1
G5
G7
EBFP
Prime editing
pegRNA
EGFP
S65T H66Y
b
EBFP
PE*
Indels
PE*
EBFP
EGFP
c
100 µM Ouabain
PE2-EBFP
100 µM Ouabain
PE3b-EBFP (G1)
PE3-ATP1A1-Q118R
99.8% 0.0% 0.2% 0.0%
83.9% 15.5% 0.1% 0.6%
49.9% 45.6% 0.1% 4.4%
78.8% 19.4% 0.1% 1.8%
30.9% 54.7% 0.2% 14.3%
EBFP
EGFP
100 µM Ouabain
PE3-EBFP (G2)
100 µM Ouabain
PE3-EBFP (G3)
100 µM Ouabain
PE3-EBFP (G4)
PE3-ATP1A1-Q118R
79.2% 16.5% 0.5% 3.9%
23.4% 40.9% 3.5% 32.1%
79.1% 12.3% 2.0% 6.6%
16.3% 15.9% 16.3% 51.5%
71.1% 16.2% 1.6% 11.1%
9.5% 14.0% 12.8% 63.7%
EBFP
EGFP
100 µM Ouabain
PE3-EBFP (G5)
100 µM Ouabain
PE3-EBFP (G6)
100 µM Ouabain
PE3-EBFP (G7)
PE3-ATP1A1-Q118R
73.1% 13.4% 5.4% 8.1%
22.7% 14.8% 25.6% 36.8%
66.5% 13.6% 4.6% 15.4%
9.9% 8.4% 14.0% 67.6%
62.2% 15.8% 4.4% 17.6%
7.6% 10.7% 10.1% 71.5%
EBFP
EGFP

**Supplementary figure 21 (Related to Fig. 6). Representative FACS plots for EBFP to EGFP conversion.** (**a**) Schematic representation of the EBFP to EGFP reporter system using different nick sgRNAs with PAMs facing outwards (PAM-Out) or inwards (PAM-In) towards the pegRNA. (**b**) Schematic of the FACS-based quantification of EBFP to EGFP conversion via PE. In K562 cells homozygous for EBFP integration at *ATP1A1* (triallelic), FACS-based analysis reports PE outcomes indirectly; (i) EBFP(+)/EGFP(+) cells result from monoallelic or biallelic PE, (ii) EBFP(-)/EGFP(+) cells are generated by either triallelic PE or combinations of mono- and biallelic PE along with indels*, (iii) EBFP(-)/EGFP(-) occur from indels on the three copies of the reporter. Indels are defined broadly in this context as any edits that inactivates the reporter. (**c**) Representative FACS plots from the EBFP to EGFP conversion experiment. As described in **Fig. 6**, K562 cells stably expressing the EBFP reporter from the *ATP1A1* locus were transfected with PE3max vectors targeting *ATP1A1* exon 4 (pegRNA-Q118R_v1) and EBFP (epegRNA). Cells were treated (ouabain) or not (untreated) with 100 µM ouabain starting 3 days post-transfection until all non-resistant cells were eliminated. *n* = 3 independent biological replicates performed at different times with equivalent results (See **Fig. 6**).

a

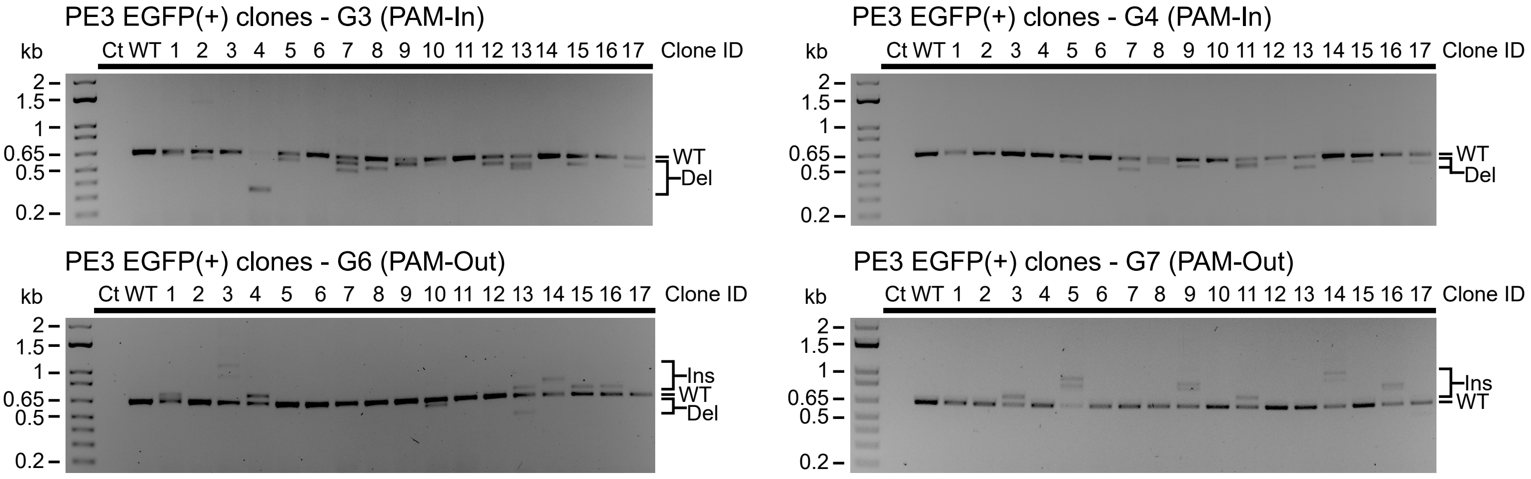
PE3 EGFP(+) clones - G3 (PAM-In)
kb
Ct WT 1 2 3 4 5 6 7 8 9 10 11 12 13 14 15 16 17 Clone ID
2
1.5
1
0.65
0.5
0.2
WT
Del
PE3 EGFP(+) clones - G4 (PAM-In)
kb
Ct WT 1 2 3 4 5 6 7 8 9 10 11 12 13 14 15 16 17 Clone ID
2
1.5
1
0.65
0.5
0.2
WT
Del
PE3 EGFP(+) clones - G6 (PAM-Out)
kb
Ct WT 1 2 3 4 5 6 7 8 9 10 11 12 13 14 15 16 17 Clone ID
2
1.5
1
0.65
0.5
0.2
Ins
WT
Del
PE3 EGFP(+) clones - G7 (PAM-Out)
kb
Ct WT 1 2 3 4 5 6 7 8 9 10 11 12 13 14 15 16 17 Clone ID
2
1.5
1
0.65
0.5
0.2
Ins
WT

**Supplementary figure 22 (Related to Fig. 6). Assessing the impact of complementary-strand nick locations on prime editing byproducts.** (**a**) PCR-based genotyping of EGFP(+) single cell-derived K562 clones targeted with a PAM-in or a PAM-out configuration. As described in **Fig. 6**, K562 cells stably expressing the EBFP reporter from the *ATP1A1* locus were transfected with PE3max vectors targeting *ATP1A1* exon 4 (pegRNA-Q118R_v1) and EBFP (epegRNA). Single cell-derived clones were isolated in methylcellulose-based semi-solid RPMI media supplemented with 100 µM ouabain and genomic DNA from clones that were EGFP(+) was harvested after coselection. Larger insertions and deletions are indicated.

**Supplementary table 1: Distribution of alleles in single cell-derived clones edited at *ATP1A1* via prime editing.** K562 cells were transfected with PE vectors targeting *ATP1A1* exon 4 (Q118R). Single cell-derived K562 clones were isolated in methylcellulose-based semi-solid RPMI media supplemented with 0.5 µM ouabain and genomic DNA was harvested after coselection. The number of precise prime edited alleles was determined from BEAT Sanger sequencing trace analysis and small insertions and deletions were analyzed with DECODR. PCR-based genotyping was performed to identify larger insertions and deletions.

| PE strategy | % of clones with indicated alleles | | | |
|---|---|---|---|---|
| | WT + Indels | Monoallelic | Biallelic | Triallelic[&] |
| PE2-Q118R[*] | 0% (0/47) | 98% (46/47) | 2% (1/47) | 0% (0/47) |
| PE3-Q118R[**] | 0% (0/47) | 45% (21/47) | 40% (19/47) | 15% (7/47) |

[&]Homozygous, only PE-specified sequences are detected

[*]No small indels or large insertions and deletions are detected, remaining alleles are WT

[**]For monoallelic and biallelic clones, remaining alleles are WT, small indels or large insertions and deletions. Scaffold incorporations are also detected on PE alleles

**Supplementary table 2: Distribution of alleles in single cell-derived clones edited at *ATP1A1* and *MTOR* by coselection using engineered pegRNAs (epegRNAs) in a PE2 mode.** K562 cells stably expressing the mScarlet-I mTOR Signaling Indicator (mSc-TOSI) reporter were transfected with PE2max-epegRNA vectors targeting *ATP1A1* exon 4 (Q118R) and *MTOR*. Single cell-derived K562 clones were isolated in methylcellulose-based semi-solid RPMI media supplemented with 100 µM ouabain and genomic DNA was harvested after coselection. The number of precise prime edited alleles was determined from BEAT Sanger sequencing trace analysis and small insertions and deletions were analyzed with DECODR. PCR-based genotyping was performed to identify larger insertions and deletions.

| Targeted locus and mutation | % of clones with indicated alleles | | | |
|---|---|---|---|---|
| | WT | Monoallelic* | Biallelic* | Triallelic& |
| *ATP1A1*-Q118R | 0% (0/20) | 10% (2/20) | 45% (9/20) | 45% (9/20) |
| *MTOR*-F2108L | 15% (3/20) | 20% (4/20) | 50% (10/20) | 15% (3/20) |

*No small indels or large insertions and deletions are detected, remaining alleles are WT

&Homozygous, only PE-specified sequences are detected
